# Structural insights into agonist binding and activation of succinate receptor 1

**DOI:** 10.1101/2024.02.03.578529

**Authors:** Aijun Liu, Yezhou Liu, Weijia Zhang, Richard D. Ye

**Affiliations:** Dongguan Songshan Lake Central Hospital, Dongguan Third People’s Hospital. Dongguan, Guangdong 523326, China; Kobilka Institute of Innovative Drug Discovery, School of Medicine, The Chinese University of Hong Kong, Shenzhen, Guangdong 518172, China; The Chinese University of Hong Kong, Shenzhen Futian Biomedical Innovation R&D Center, Shenzhen, Guangdong 518000, China

## Abstract

Succinate is an intermediate of the citric acid cycle and serves important functions in energy homeostasis and metabolic regulation. Extracellular accumulation of succinate acts as a stress-induced signal through its G protein-coupled receptor, SUCNR1. Research on succinate signaling is hampered by the lack of high-resolution structures of the agonist-bound receptor. Here we present cryo-EM structures of SUCNR1-Gi complexes with the receptor bound to succinate and its non-metabolite derivative epoxysuccinate. Structural analysis of SUCNR1 identified key determinants for recognition of the dicarboxylate agonists in *cis* conformation. R281^7.39^ and Y83^2.64^ are critical to ligand binding, but Y30^1.39^ and R99^3.29^ also participate in binding of succinate and epoxysuccinate, respectively. The extracellular loop 2, through F175^ECL2^ in its β-hairpin, forms a hydrogen bond with one of the carboxyl groups and serves as a lid to cap the binding pocket for succinate. At the receptor-Gi protein interface, agonist binding induces the rearrangement of a hydrophobic network on TM5 and TM6, leading to transmembrane signaling through TM3 and TM7. The agonist-bound SUCNR1 structures shed light on molecular recognition of succinate for receptor signaling, that may promote further development of novel agonists, antagonists and biased agonists targeting SUCNR1.

## Introduction

Succinate is a small dicarboxylic acid produced as an intermediate of the tricarboxylic acid cycle. Succinate also serves as a substrate of the mitochondrial respiratory chain through oxidation by succinate dehydrogenase (SDH). Under conditions such as hypoxia (Chouchani et al., 2014), inflammation (Keiran et al., 2019; Peruzzotti-Jametti et al., 2018), infection (Perniss et al., 2023), and tissue damage (Hamel et al., 2014; Sapieha et al., 2008), succinate is secreted extracellularly as autocrine and paracrine signals. Extracellular succinate is a pleiotropic hormone-like metabolite that acts via its receptor, succinate receptor 1 (SUCNR1) (Fernandez-Veledo et al., 2021; He et al., 2004), that is widely expressed in human tissues (Gilissen et al., 2016) including epithelia of the intestine and kidney (Schneider et al., 2018), white adipose tissue and immunological tissue (Rubic et al., 2008). The close relationship between SUCNR1 and many inflammatory and metabolic diseases such as liver fibrosis (Winther et al., 2021), type 2 diabetes (Villanueva-Carmona et al., 2023), rheumatoid arthritis (Littlewood-Evans et al., 2016), dermatitis (Gnana-Prakasam et al., 2011), cancer metastasis (Wu et al., 2020), obesity (Keiran et al., 2019), and hypertension (Sadagopan et al., 2007), make this receptor an attractive target for therapeutic intervention.

The SUCNR1 gene was located on human chromosome 3q24-3q25 and was first identified to encode an orphan G protein-coupled receptor (GPCR) named GPR91 (Gonzalez et al., 2004; Wittenberger et al., 2001). It was speculated to be a nucleotide receptor due to its high sequence homology with purinergic receptors (Abbracchio et al., 2006). In a landmark study employing mass spectrometry and functional assays, the natural ligand of SUCNR1 was found to be succinate (He et al., 2004). The binding of succinate to SUCNR1 triggers primarily Gi protein-coupled pathways, with half maximal effective concentration (EC_50_) of 17-56 μM (He et al., 2004). Effort has been made to improve agonist stability and potency, resulting in a synthetic succinate analogue, *cis*-epoxysuccinate, as a full agonist with 10-fold higher potency than succinate (EC_50_ = 2.7 μM) (Trauelsen et al., 2017). *Cis*- epoxysuccinate belongs to one of the most potent non-metabolite synthetic agonists of SUCNR1, which is not a substrate of succinate dehydrogenase (SDH) (Trauelsen et al., 2017).

Despite emerging interests in SUCNR1 as a potential drug target, the structural basis for succinate recognition and G protein activation remains unclear. Several laboratories have used computer modelling to predict succinate binding pocket in SUCNR1 (Geubelle et al., 2017; He et al., 2004; Trauelsen et al., 2017). Since these models were built on the structurally homologous P2Y1 receptor and focused primarily on the positively charged arginines for binding to the negatively charged bicarboxylates in succinate, there were significant discrepancies. A crystal structure of rat SUCNR1 in apo state was obtained by Haffke and coworkers (Haffke et al., 2019). Another structure with the humanized rat SUCNR1 bound to an antagonist NF-56-EJ40 was also reported in the same paper (Haffke et al., 2019). These structures were obtained with the use of nanobody6 which stabilizes the receptor structure, without the addition of succinate to the receptor preparation. Therefore, although the study showed the overall structure of SUCNR1, it did not address the questions of succinate interaction with SUCNR1 and succinate-induced G protein activation. To better comprehend the ligand recognition and activation mechanism, we purified the human SUCNR1-Gi protein complexes bound to the natural agonist succinate and to a synthetic non-metabolite agonist *cis*- epoxysuccinate. The structures of these protein complexes were determined by single particle cryo-electron microscopy (cryo-EM). Combined with molecular dynamics simulation and mutagenesis assays, these structures provide insights into the G protein coupling mechanism as well as ligand recognition. The molecular details may enhance our understanding of dicarboxylic acid recognition by GPCR, and promote further development of agents that target SUCNR1.

## Methods

### Cloning and purification of the SUCNR1- Gαi1 complex

The coding sequence of human SUCNR1 was cloned into a pFastBac1 vector for expression in *Sf*9 insect cells. To facilitate Protein purification and improve protein thermal stability, an N terminal hemagglutinin (HA) signal peptide, a FLAG-tag, a human rhinovirus 3C (HRV 3C) protease cleavage site (LEVLFQGP) and thermostabilized apocytochrome b562RIL (BRIL) weren fused. The widely used NanoBiT tethering strategy was introduced for structure determination as reported (Chen et al., 2022; Dixon et al., 2016; Duan et al., 2020; You et al., 2023) . The human Gαi1 with two dominant-negative mutations (G203A, A326S) cloned into the pFastBac1 vector, Gβ1 and Gγ2 cloned into the pFastBac-Dual vector was used as described previously (Wang et al., 2023). For functional assays, the human SUCNR1 coding sequence was cloned into the pcDNA3.1 vector. Point mutations were generated by PCR-mediated site-directed mutagenesis.

The baculovirus expression system was used for protein expression. Baculoviruses of SUCNR1, Gαi1 and Gβ1γ2 were generated according to the Bac-to-Bac manual (Thermo Fisher Scientific) and transfected into Sf9 cells at a density of 2×10^6^ for co-expression. After 48h of transfection, the cell culture was harvested by centrifugation at 2000 x g for 15 min and kept frozen at −80 °C until use.

For purification of the succinate-SUCNR1-Gi and epoxysuccinate-SUCNR1-Gi complex, cell pellets were resuspended in the buffer containing 25 mM HEPES pH 7.4, 50 mM of NaCl, 5 mM of KCl, 5 mM of MgCl_2_, 5 mM of CaCl_2_, 5% glycerol, 25 mU/mL apyrase, 2.5 μg/ml leupeptin, 0.16 mg/ml benzamidine, 100 μM of agonists (succinate or *cis*-epoxysuccinate) were added before incubation for 30 min. Cell membranes were collected by centrifugation and solubilized in 20 mM HEPES pH 7.4, 100 mM NaCl, 1% LMNG, 0.1% CHS, 10% glycerol, 2.5 μg/ml leupeptin, 0.16 mg/ml benzamidine and 50 M of agonists. The supernatant was cleared by centrifugation at 20,000 × g for 35 mins and loaded onto a gravity-flow column to incubate with anti-FLAG affinity resin (GenScript Biotech). The resin was washed with 15 column volumes of Wash Buffer containing 20 mM of HEPES (pH 7.4), 100 mM of NaCl, 5% glycerol, 2 mM of MgCl_2_, 2 mM of CaCl_2_, 0.05% (w/v) LMNG, 0.05% (w/v) GDN (Anatrace), 0.003% (w/v) CHS, and 20 M of agonists. The protein complex was then eluted with buffer containing 20 mM of HEPES (pH 7.4), 100 mM of NaCl, 2 mM of MgCl_2_, 2 mM of CaCl_2_, 0.01% (w/v) LMNG, 0.001% (w/v) GDN (Anatrace), 0.001% (w/v) CHS, 0.2 mg/ml FLAG peptide (Sigma-Aldrich), and 20 μM of the agonists. The eluted fraction was concentrated using an Amicon® Ultra-15 Centrifugal Filter Unit (Millipore) and subjected to size exclusion chromatography on tandem Superose 6 10/300 and Superdex 200 Increase 10/300 column (Cytiva) pre-equilibrated with 20 mM HEPES (pH 7.4), 100 mM NaCl, 0.0015% LMNG, and 0.0005% GDN, 0.0003% CHS and 20 μM of agonists. The peak fractions corresponding to the protein complex were collected, analyzed on SDS-PAGE and Western-blotting, concentrated to approximately 10 mg/mL, and stored at −80°C until further use.

### Cryo-grid preparation and EM data collection

Before cryo-grid preparation, negative stain electron microscopy was performed on all the samples to confirm homogeneity and complex formation. Aliquots of 3uL of purified protein complex were applied onto a glow-discharged Ultrafoil 300 mesh R1.2/1.3 holy Au grid (Tergeo-EM plasma cleaner). The grids were blotted for 3.5 s with a blot force of 1 in 100% humidity at 4 °C, and then quickly plunged into liquid ethane using a Vitrobot Mark IV (Thermo Fisher Scientific).

The grid sample screening and data collection were performed by SerialEM software (Mastronarde, 2005) installed on a 300 kV Titan Krios Gi3 microscope. The final data sets were collected with a Gatan K3 direct electron detector at a nominal magnification of 105,000 (calibrated pixel size of 0.85 Å). The movie stacks were acquired with a total exposure time of 2.5s fractionated to 50 frames and a dose rate of 21.3 e/pixel/s. The defocus range was set from −1.2 to −2.5 μm and a GIF Quantum energy filter (Gatan, USA) was used to exclude inelastically scattered electrons with a slit width of 20 eV.

### Image processing and 3D reconstructions

The overall cryo-EM datasets were processed by cryoSPARC version v4.2.1 (Punjani et al., 2017) and Relion version 4.0 (Kimanius et al., 2021). All movie stacks were aligned with motion correction and dose-weighting. After contrast transfer function (CTF) estimation, micrographs were manually inspected and obvious bad micrographs were discarded. Initial two-dimensional (2D) templates for autopicking were generated by 2D classification of manually picked particles.

For the succinate-SUCNR1-Gi dataset, a total of 2,043,885 particles were template-based picked. The particles were then subjected to 3 rounds of 2D classification and particles in 2D averages with clear secondary features were selected. Ab initio reconstruction was performed by cryoSPARC followed by rounds of 3D classification. Finally, a dataset of 147371 particles was subjected to homogeneous refinement, non-uniform refinement, and local refinement. The global resolutions estimated by the ‘gold standard’ criterion (FSC = 0.143) was 2.97 Å.

For the epoxysuccinate-SUCNR1-Gi dataset, template-based particle picking resulted in a dataset containing 2024614 particles. After multiple rounds of classification and manual selection, a total of 271253 particles yielded a final map with an estimated global resolution of 3.15 Å.

### Model building and refinement

The model building was facilitated by the previous structure of PDB code 6IBB (Haffke et al., 2019) and 8JJP (Liu et al., 2023), as a starting template for SUCNR1 and G protein respectively. The model was manually adjusted and built by Coot (Emsley et al., 2010), as well as iterative real-space refinement in Phenix (Liebschner et al., 2019). The final model validations were carried out by Molprobity (Chen et al., 2010). The molecular graphic figures were prepared by UCSF Chimera (Pettersen et al., 2004), ChimeraX (Pettersen et al., 2021) and PyMoL software.

### G protein dissociation assay

G protein activation was assessed through a NanoBiT-based G protein dissociation assay (Inoue et al., 2019). HEK293T cells were seeded in a 24-well plate 24 hours prior to transfection. Transfection with Lipofectamine™ 3000 (Invitrogen, L3000001) involved a mixture of 92 ng pcDNA3.1 vector encoding human SUCNR1 (wild type/mutants), 46 ng pcDNA3.1 vector encoding Gαi1-LgBiT, 230 ng pcDNA3.1 vector encoding Gβ1, and 230 ng pcDNA3.1 vector encoding SmBiT-Gγ2 (per well in a 24-well plate). After 24 hours of incubation, the transfected cells were collected and suspended in HBSS containing 20 mM HEPES. 20 μL cell suspension was placed on a 384-well white plate (PerkinElmer Life Sciences, Waltham, MA) and loaded with 5 μL of 50 μM coelenterazine H (Yeasen Biotech, Shanghai, China). Following a 2-hour incubation at room temperature, the baseline was measured using an Envision 2105 multimode plate reader (PerkinElmer). Subsequently, either succinate (A610496, Sangon Biotech, Shanghai, China) or *cis*-epoxysuccinate (HY-125791, MCE, Monmouth Junction, NJ) were applied to the cells at varying concentrations. The luminescence signals induced were measured 15 minutes after ligand addition and normalized by dividing them by the initial baseline readings. The fold changes in signals were further normalized to the signal from HBSS-treated negative control samples, and the EC_50_ values were calculated relative to different ligand concentrations based on three independent experiments, each with triplicate measurements.

### cAMP assay

Human SUCNR1, both in its wild-type form and in mutants, were expressed in HeLa cells 24 hours before harvesting. The cells were suspended in HBSS, containing 5 mM HEPES, 0.1% BSA (w/v), and 0.5 mM 3-isobutyl-1-methylxanthine, then loaded onto 384-well plates. Different concentrations of succinate or epoxysuccinate ligands were prepared along with 2.5 μM forskolin in the above-mentioned buffer were prepared. Following this, the cells were stimulated with the ligands and 2.5 μM forskolin for 30 minutes in a cell incubator. Measurement of intracellular cAMP levels was carried out using the LANCE Ultra cAMP kit (PerkinElmer, TRF0263), adhering to the manufacturer’s instructions. During the measurements, signals from time-resolved fluorescence resonance energy transfer (TR-FRET) were recorded using an EnVision 2105 multimode plate reader (PerkinElmer). The determination of intracellular cAMP levels was based on the TR-FRET signals of the samples in comparison to cAMP standards.

### Flow cytometry analysis

HEK293T cells underwent transfection with expression plasmids encoding FLAG-tagged wild-type (WT) or mutant SUCNR1 for 24 hours at 37°C. Subsequently, the cells were collected and washed with HBSS containing 0.5% BSA. Following this, the cells were incubated with a FITC-labeled anti-FLAG antibody (Sigma, Cat #F4049; diluted 1:50 in HBSS buffer) for 30 minutes on ice and then washed with HBSS. Flow cytometry (CytoFLEX, Beckman Coulter) was employed to quantify the FITC fluorescence signals, indicative of the antibody-receptor complex on the cell surface. The fluorescence signals were assessed for the relative expression levels of SUCNR1 mutants.

### Molecular dynamics (MD) simulation

MD simulations were conducted using Gromacs-2020.4 (Van Der Spoel et al., 2005). The missing residues due to internal flexibility in the N terminal were modeled through the MODELLER program (Webb & Sali, 2016) based on prediction from AlphaFold2 (Jumper et al., 2021). The protonation state of charged residues was assigned by assuming pH 7.4. The N and C termini were end-capped with acetyl or N-methyl groups for amines or carboxylic acids, respectively. The ligand-protein complex was embedded into a POPC bilayer membrane by CHARMM-GUI membrane builder (Wu et al., 2014) with the orientation estimated by PPM2.0 (Lomize et al., 2012). The TIP3P model was used for water, and the system was solvated in a water box and charge-neutralized by 150 mM NaCl. The CHARMM36m force field (Huang et al., 2017) was used for simulation, and the parameters of ligands were generated by the CgenFF program (Vanommeslaeghe & MacKerell, 2012; Vanommeslaeghe et al., 2012).

The inputs for Gromacs were prepared using CHARMM-GUI (Wu et al., 2014). After energy-minimization and pre-equilibration, three independent 400-ns long product simulations were performed with temperature and pressure set at 310.15 K and 1 bar by Nose-Hoover thermostat and Parrinello-Rahman barostat respectively. Long-range electrostatic interactions were calculated by the particle mesh Ewald (PME) method with a cutoff of 12 Å. The Van der Waals interactions were cut off by smoothly switching to zero starting at 10 Å to 12 Å. All bonds were constrained using the Linear Constraint Solver (LINCS) algorithm.

### Statistical Analysis

The analysis of the data was conducted using Prism 9.5.0 (GraphPad, San Diego, CA). Dose-response curves for agonist analysis were generated employing the log[agonist] vs. response equation (three parameters) within the software. In the case of cAMP and G protein dissociation assays, data points were expressed as percentages (mean ± SEM) relative to the maximal response level for each sample, derived from a minimum of three independent experiments, as specified in figure legends. The EC_50_ values were extracted from the dose-response curves. Regarding cell surface expression, data points were displayed as percentages (mean ± SEM) of the flow cytometry fluorescence signals of wild-type SUCNR1. Statistical comparisons were performed using Analysis of Variance (ANOVA) through the one-way method. A *p*-value of 0.05 or lower was deemed statistically significant.

## Results

### Overall structures of SUCNR1 signaling complexes

To solve the structure of SUCNR1, an N-terminal FLAG-tagged human full-length SUCNR1 and heterotrimeric G proteins (DNGi, Gβγ) were co-expressed in *Sf*9 insect cells. The natural agonist succinate and synthetic non-metabolic agonist *cis*- epoxysuccinate of SUCNR1 were added in respective samples during protein purification to facilitate the formation of signaling complexes. The protein complexes were purified by tandem affinity chromatography (anti-FLAG) and size exclusion chromatography (SEC; Fig. S1, S2). The homogeneity of the protein complexes was further evaluated by transmission electron microscopy (Fig. S1, S2) and the structures were determined by cryo-electron microscopy (cryo-EM) single particle analysis, obtaining the succinate-SUCNR1-Gi complex at a nominal global resolution of 2.97 Å (Fig. 1A) and the epoxysuccinate-SUCNR1-Gi complex at 3.15 Å (Fig. 1B). The density of the ligand was well distinguished, and the density of the sidechains of most residues was clearly defined. The quality of the EM map enabled us to unambiguously build molecular models of the succinate-SUCNR1-Gi complex (Fig. 1C, 1E; Fig. S3) and the epoxysuccinate-SUCNR1-Gi complex (Fig. 1D, 1F; Fig. S4) except for their disordered termini. The description and statistics are presented in Table S1 and *Methods*. The overall arrangement of the two SUCNR1 signaling complexes are highly similar and largely resemblant to that of other Class A GPCRs, including a canonical seven transmembrane (TM) bundle architecture and an intracellular amphipathic helix. The seven transmembrane helices surround the central orthosteric ligand-binding pocket on the extracellular side and form close contact with the G protein on the intracellular side. There are two disulfide bridges in SUCNR1, all formed on the extracellular side. One is formed between C11^1.20^ and C268^7.26^ [Ballesteros–Weinstein numbering for GPCRs (Ballesteros & Weinstein, 1995) is present as superscripts] that stabilizes the N terminus and the TM7 (Haffke et al., 2019) . The second one is between C95^3.25^ and C172^ECL2^, found in almost all Class A GPCRs and plays an important role in GPCR function (Fernandez-Veledo et al., 2021) . Notably, there was an additional β-strand and a small hairpin in the second extracellular loop (ECL2; Fig. 1E, 1F), which was not found in the inactive SUCNR1 structure (Haffke, 2019). The β-hairpin in ECL2 is similar to the one found in the structure of activated P2Y1 (Li et al., 2023) except that the one in this study is more vertical. The small helix in ECL2 closer to the agonist binding pocket may form an upper lid to prevent release of the bound ligand (Wheatley et al., 2012). Compared with P2Y1, the binding pocket of succinate is deeper and aligned with amino acid side chains that form hydrophobic interaction with the ligand in addition to direct polar interactions (Fig. S5, S6).

**Fig. 1.**
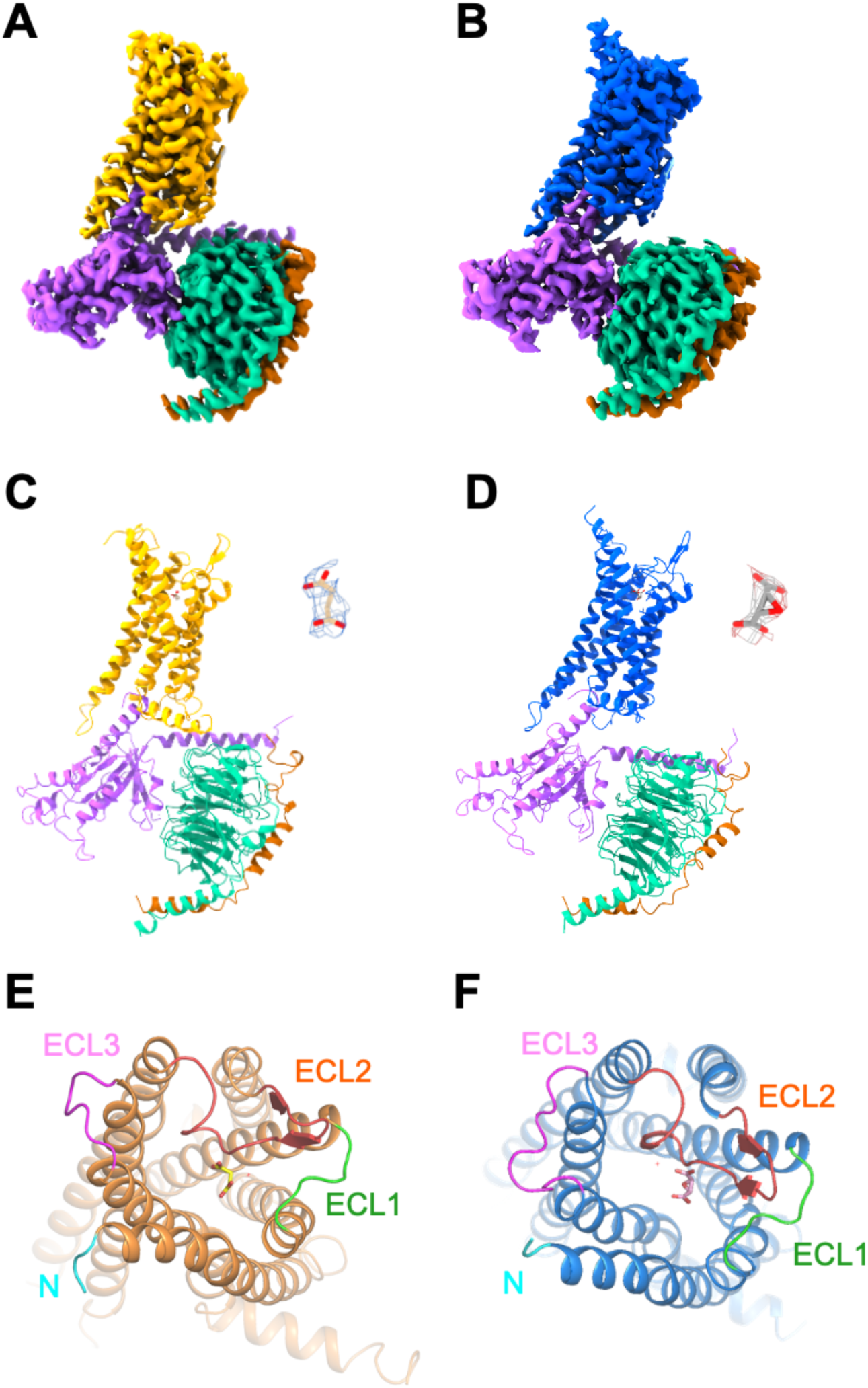
Structure of the SUCNR1-Gi bound to succinate and epoxysuccinate. **(A)** Cryo-EM density map of the succinate-bound SUCNR1-Gi complex. **(B)** Cryo-EM density map of the epoxysuccinate-bound SUCNR1-Gi complex **(C)** Overall structure of succinate-bound SUCNR1-Gi complex (side view). Structure and EM density of succinate is highlighted on the right. **(D)** Overall structure of epoxysuccinate-bound SUCNR1-Gi complex (side view). Structure and EM density of epoxysuccinate is highlighted on the right. **(E)** Structure of succinate-bound SUCNR1 from the top view. Extracellular domains are highlighted. **(F)** Structure of epoxysuccinate-bound SUCNR1 from the top view. Extracellular domains are highlighted.

### Molecular basis for succinate recognition by SUCNR1

Succinate is the natural ligand of SUCNR1. In the succinate-SUCNR1-Gi complex, succinate inserts deep into the hydrophilic and electropositive transmembrane pocket surrounded by TM1, TM2, TM3, TM7, and ECL2 (Fig. 2A, Fig.S7A-S7C). As a dicarboxylic acid receptor, the most notable feature of SUCNR1 is the interaction between carboxyl groups of the ligand and the receptor binding pocket. In the molecule model based on our resolved structure of the succinate-SUCNR1-Gi complex, succinate adopts a rather horizontal pose in *cis* conformation in the ligand binding pocket (Fig. 2A, 2B), with its two carboxyl groups pointing to the same extracellular direction. One of the carboxyl groups of the ligand is very close to TM1, TM2 and TM7, forming a salt bridge with R281^7.39^ and a hydrogen bond with the hydroxyl group of Y83^2.64^. This carboxyl group forms an additional weak hydrogen bond with Y30^1.39^, which also contributes to the polar interaction between the receptor and the ligand (Fig. 2B, 2C). The other carboxyl group is closer to TM3 and ECL2 of the receptor, forming a weak hydrogen bond with the main chain of F175^ECL2^. R99^3.29^ and H103^3.33^, mentioned in a prior model for interaction with succinate (He et al., 2004), are not critical in our model since their sidechains do not form direct interactions with the ligand (Fig. 2D). R252^6.55^, predicted in previous models to interact with one of the carboxyl groups (Geubelle et al., 2017; He et al., 2004), is too far away to interact with the ligand in our model (Fig. 2B, 2C).

**Fig. 2.**
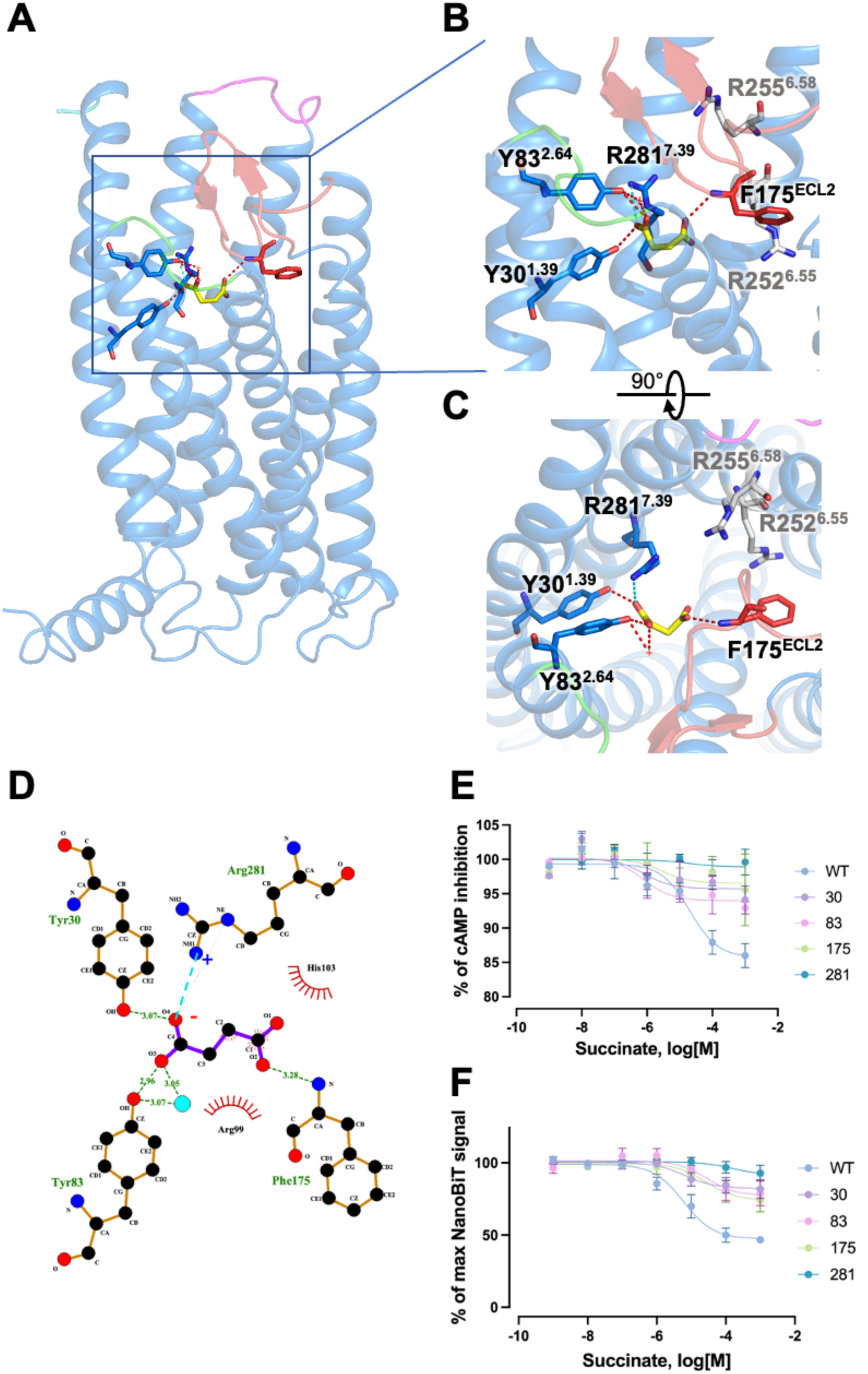
Ligand binding pocket in succinate-SUCNR1-Gi complex. **(A)** Interaction between succinate and SUCNR1. Succinate is shown in yellow sticks. β-hairpin of ECL2 is shown in red. **(B)** Polar interactions between succinate and SUCNR1 (side view). Hydrogen bonds are shown in red dashes. Salt bridges are shown in cyan dashes. R252^6.55^ and R255^6.58^, that are not engaged in succinate binding, in this model, are colored in grey sticks. **(C)** Polar interactions between succinate and SUCNR1 (top view). **(D)** 2D representation of succinate binding to SUCNR1. Hydrogen bonds are shown in green dashes. Salt bridges are shown in cyan dashes, positive and negative charged atoms are marked, respectively. **(E)** Effect of alanine substitution of selected amino acids with polar interactions to succinate in the receptor binding pocket on cAMP inhibition. **(F)** Effect on G protein dissociation of the selected SUCNR1 mutants treated with different concentrations of succinate. Data shown are means ± SEM of three independent experiments.

To functionally verify the SUCNR1 residues engaged in succinate recognition and downstream G protein signaling, we substituted the succinate-interacting residues by alanine, and conducted cAMP inhibition assay and G protein dissociation assays with the resulting mutants. Our results indicate that substitution of R281^7.39^ with alanine completely eliminated the ability of the receptor to activate Gi protein, while alanine substitutions of Y30^1.39^, Y83^2.64^ and F175^ECL2^ produced moderate reduction of signal transduction by the mutants (Fig. 2E). G protein dissociation assay confirmed these findings (Fig. 2F).

### Molecular basis for epoxysuccinate recognition by SUCNR1

Epoxysuccinate is the non-metabolic analogue of succinate with similar functions (Geubelle, 2017). An analysis of the molecular model based on our cryo-EM structure of epoxysuccinate-SUCNR1-Gi complex found that the binding pocket for epoxysuccinate is nearly identical to that for succinate, but is almost sealed by residues close to the extracellular space including Y83^2.64^, R99^3.29^, R281^7.39^ and D174^ECL2^ (Fig. 3A, Fig. S7D-S7F), creating a more isolated environment for epoxysuccinate binding. Like succinate, epoxysuccinate adopts a *cis* conformation in the SUCNR1 binding pocket. While the planar epoxy group lies horizontally at the bottom of the binding pocket, the two carboxyl groups are pointing upwards (Fig. 3B). One of the carboxyl groups interacts with R281^7.39^ through a salt bridge and forms a hydrogen bond with the hydroxyl group of Y83^2.64^. For the other carboxyl group, the interaction profile is different from succinate bound to SUCNR1. Of note, R99^3.29^ forms a salt bridge with epoxysuccinate at this carboxyl group (Fig. 3B, 3C). Moreover, Y248^6.51^ and Y277^7.35^ also participate in the interaction with the carboxyl oxygen via an intermediate water (Fig. 3D). Y277^7.35^ and D174^ECL2^ was shown in a previous report to form a hydrogen bond (Haffke et al., 2019). This bond was broken in our model, allowing the formation of a new salt bridge between R281^7.39^ and D174^ECL2^ that further stabilizes the active conformation of the receptor (Fig. S8). The interactions between epoxysuccinate and SUCNR1 are further analyzed through MD simulations, and results indicate that the salt bridges formed between the ligand and R99^3.29^ and R281^7.39^, respectively, as well as the hydrogen bond between the ligand and Y83^2.64^ are stable throughout the 3×400 ns MD simulation (Fig. S9).

**Fig. 3.**
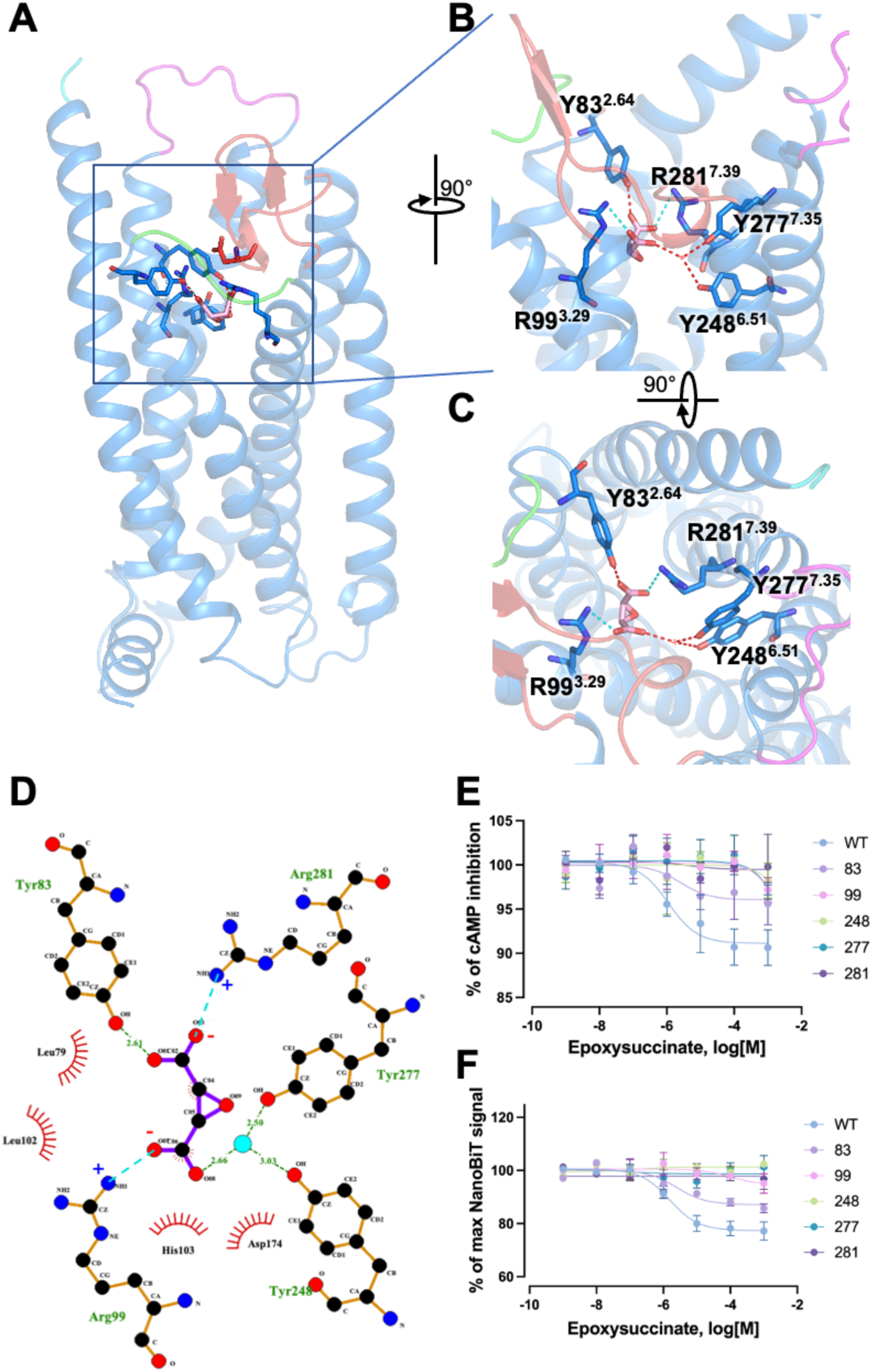
Ligand binding in epoxysuccinate-SUCNR1-Gi complex. **(A)** Interaction between epoxysuccinate and SUCNR1. Epoxysuccinate is shown in pink sticks. **(B)** Polar interactions between epoxysuccinate and SUCNR1 (side view, turned 90° from Panel A for better visualization). Hydrogen bonds are shown in red dashes. Salt bridges are shown in cyan dashes. **(C)** Polar interactions between epoxysuccinate and SUCNR1 (top view). **(D)** 2D representation of epoxysuccinate binding to SUCNR1. Hydrogen bonds are shown in green dashes. Salt bridges are shown in cyan dashes, positive and negative charged atoms are marked, respectively. **(E)** Effect of alanine substitution of selected amino acids with polar interactions to epoxysuccinate in the receptor binding pocket on cAMP inhibition. **(F)** Effect on G protein dissociation of the selected SUCNR1 mutants treated with different concentrations of epoxysuccinate. Data shown are means ± SEM of three independent experiments.

Alanine substitution of epoxysuccinate binding residues was performed, along with G protein signaling assays upon epoxysuccinate ligand treatment. The surface expression levels of all aforementioned mutants were analyzed by flow cytometry, and comparable to the expression level of wild type SUCNR1 (Fig. S10). In cAMP reduction assay that reflects Gi activation, alanine substitution of R281^7.39^ abrogated cAMP reduction mediated by the resulting receptor. Alanine substitutions of Y277^7.35^, Y248^6.51^ and R99^3.29^ produced intermediate effects in SUCNR1 signaling, with a smaller effect found in the Y83^2.64^A mutation (Fig. 3E). Likewise, G protein dissociation assay agreed with the above results (Fig. 3F). These findings support the respective roles of the amino acids in their interactions with epoxysuccinate as predicted by our model.

### The G protein interface of SUCNR1

In the SUCNR1-Gi complex, the Gi protein coupled to the receptor in a canonical way but with some distinct features. Compared to several Gi-coupled receptors including P2Y1 (7XXH), HCAR2 (8IHB), CB1 (6N4B) and NTSR (6OS9), the α5 helix is more vertical, and the αN helix and Ras-like domain are further away from the ICL2 and ICL3 of SUCNR1, respectively (Fig. 4A). The α5 helix in Gαi packs closely against TM6 and dominates the interaction between Gi and SUCNR1 (Fig. 4B). The intracellular pocket entangling the Gαi protein is formed mainly by TM3, TM5, TM6, ICL2, and ICL3. Hydrophobic interaction constitutes the major component of interaction between the Gαi subunit and SUCNR1, although a salt bridge between R217^5.68^ and D341^G.H5.13^, and a weak hydrogen bond between the main chain of P127^ICL2^ and N347^G.H5.19^ in the succinate-SUCNR1-Gαi structure are also observed (Fig. 4C). The conserved residues A224^6.27^, L225^6.28^, L233^6.36^, L227^6.30^, P230^6.33^, I123^3.53^, P127^ICL2^ and F128^ICL2^ of SUCNR1 form a hydrophobic network, interacting with I343^G.H5.15^, I344^G.H5.16^, L348^G.H5.20^, L353^G.H5.25^ in the α5 helix of Gαi subunit (Fig. 4C).

**Fig. 4.**
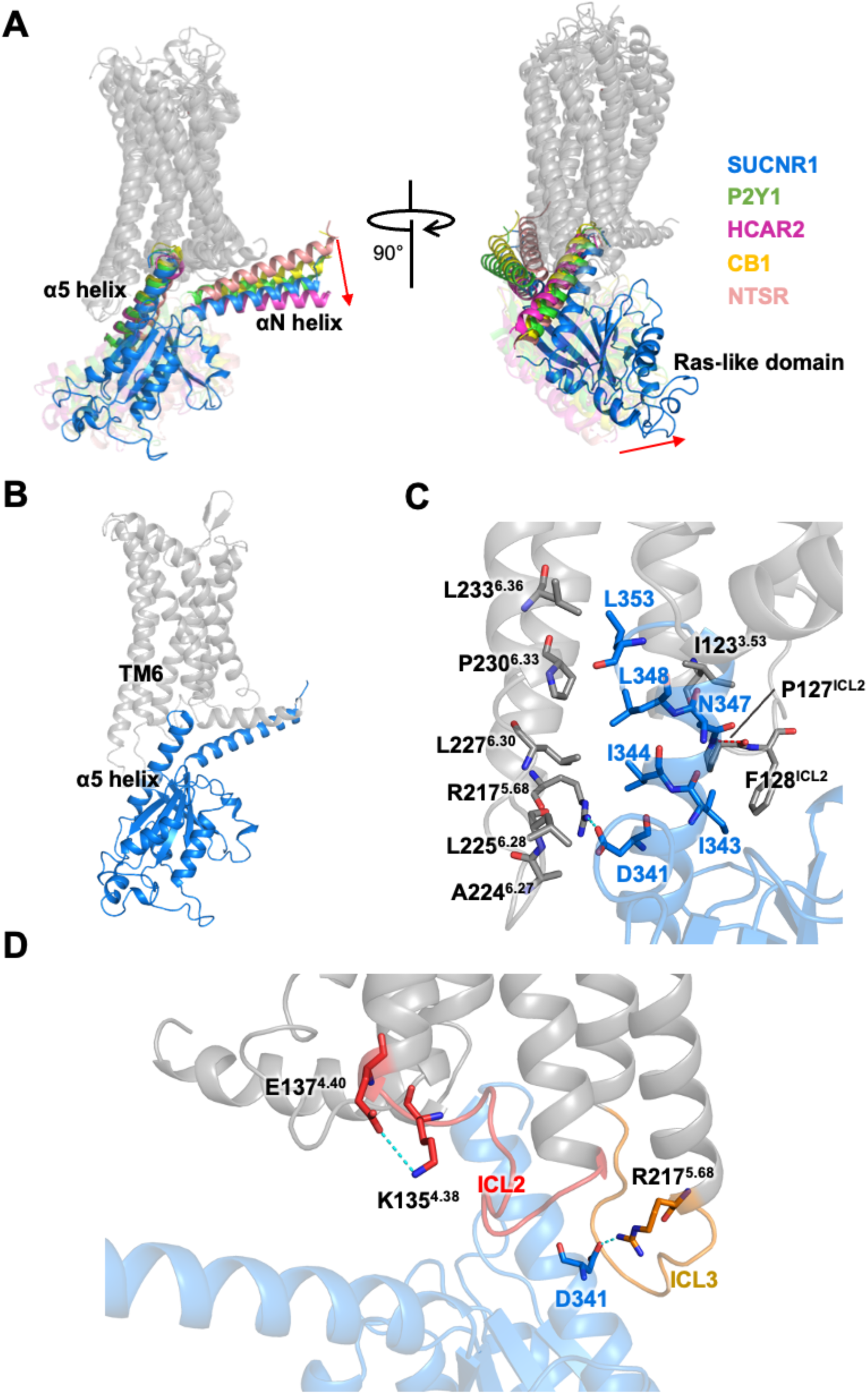
The G protein interfaces of succinate-bound SUCNR1-Gi complex. **(A)** Comparison of G protein interfaces among succinate-bound SUCNR1 (Gα shown in marine blue), P2Y1 (PDB ID: 7XXH; α5 helix and α5 helix in Gαi shown in green), HCAR2 (PDB ID: 8IHB; α5 helix and α5 helix in Gαi shown in magenta), CB1 (PDB ID: 6N4B; α5 helix and α5 helix in Gαi shown in yellow), NTSR (PDB ID:6OS9; α5 helix and α5 helix in Gαi shown in salmon). Receptors are shown in grey and aligned with each other. **(B)** The interaction between Gi and SUCNR1 is dominated by the packing between α5 helix in Gαi and TM6 of SUCNR1. **(C)** Enlarged view showing molecular interactions between Gαi and SUCNR1. Hydrogen bond between N347 and F128^ECL2^ is shown in red dash, and the salt bridge between D341 and R217^5.68^ is shown in cyan dash. **(D)** ICL2 and ICL3 pack against the Gαi protein. ICL2 is shown in red and ICL3 is shown in orange. Residues with salt bridges are shown in stick and salt bridges are shown in cyan dashes.

In the SUCNR1-Gi interface, the ICL2 and ICL3 adopt a loop-like conformation, packing against the pocket in the Gαi subunit (Fig. 4D). The salt bridges formed by D119^3.49^ (D119^3.49^-R129^ICL2^ and D119^3.49^-R120^3.50^) in the inactive structure (Haffke et al., 2019) are broken and replaced by a new salt bridge near ICL2, between K135^4.38^ and E137^4.40^, that stabilizes the active conformation of SUCNR1 (Fig. 4D). Subtle differences exist in the conformation of the ICL3 between the succinate-bound SUCNR1 structure and the epoxysuccinate-bound structure, with the former having an additional salt bridge between R217^5.68^ and D341^G.H5.13^(Fig. 4D, Fig. S11). The functional impact of this subtle difference is presently unknown.

### Structural basis for agonist-induced activation of SUCNR1

Based on our models, the succinate- and epoxysuccinate-bound SUCNR1-Gi complexes display features common to activated GPCRs (Fig. 5). To allow the coupling of Gi to SUCNR1, the TM5 and TM6 move outward, and the TM7 moves inward. Specifically, the ICL4 that connects TM7 and Helix 8 is also moved outward for accommodation of α5 helix of Gαi (Fig. 5A). There is a salt bridge between H301^8.49^ and D304^8.52^ in the inactive structure of SUCNR1 (Haffke et al., 2019), which is broken in our model for a new salt bridge between D300^8.48^ and R303^8.51^ that further stabilizes the conformation of Helix 8 (Fig. 5B). SUCNR1 contains conserved elements for GPCR activation, including DRY, PIF, and NPxxY motifs (Fig. 5C-5E). However, SUCNR1 lacks the toggle switch residue W^6.48^ of the CWxP motif with an F^6.48^ substitution, which is commonly present in the δ-branch GPCRs (Zhou et al., 2019). Likewise, general features observed in δ-branch GPCR activation, such as upward shift of TM3, are also observed in the activated SUCNR1.

**Fig. 5.**
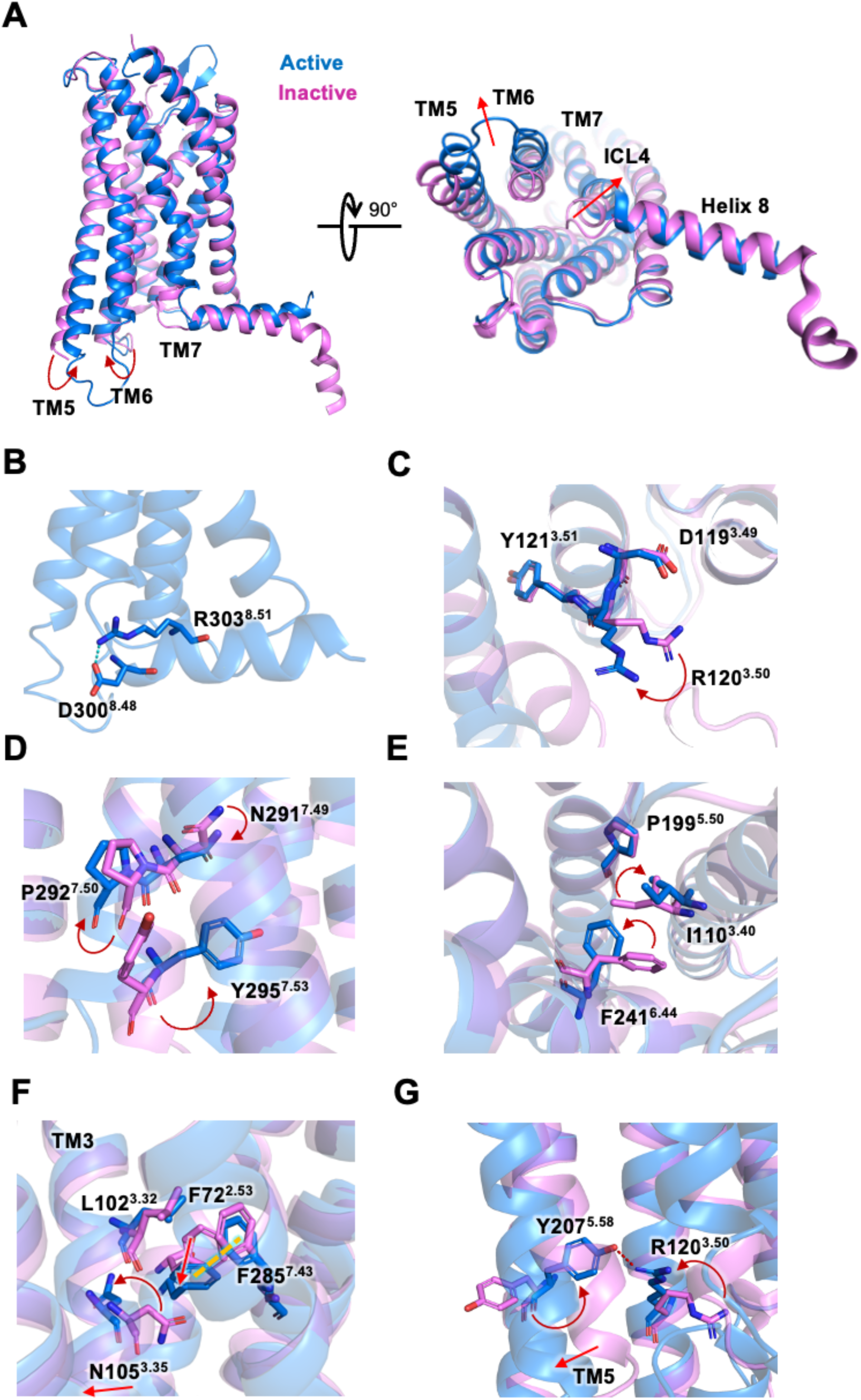
Comparison of GPCR structural motifs for receptor activation. **(A)** Conformation changes of SUCNR1 from an inactive state (PDB ID: 6IBB, shown in violet) to an active state (succinate-bound, marine blue). Conformational changes are marked with arrows. **(B)** The salt bridge between D300^8.48^ and R303^8.51^ (cyan dashes) stabilizes the conformation of Helix 8. **(C)** Close-up view of the D^3.49^-R^3.50^-Y^3.51^ motif. A clockwise turn of R^3.50^ of SUCNR1 is highlighted by a red arrow. **(D)** Close-up view of the N^7.49^P^7.50^xxY^7.53^ motif. Rotamer conformational changes of residue sidechains are shown by red arrows. **(E)** Rotamer conformational changes at the P^5.50^-I^3.40^-F^6.44^ motif. Conformational changes of residue sidechains are highlighted by red arrows. **(F)** Rearrangement of hydrophobic network in TM3. π-π stacking is shown in yellow dashes. Rotamer conformational changes of residue sidechains are highlighted in red arrows. **(G)** Rearrangement of hydrophobic network in TM5. Hydrogen bonds are highlighted in red dashes. Residue sidechain conformational changes are shown in red arrows.

Next, the structure of agonist-bound SUCNR1 is compared with an available structure of SUCNR1 bound to an antagonist, NF-56-EJ40 (6RNK; (Haffke et al., 2019)). NF-56-EJ40 is bulkier than succinate and epoxysuccinate, and therefore has more contact area in binding SUCNR1 (Fig. S12). NF-56-EJ40 differs from succinate and epoxysuccinate in that it carries only one carboxyl group and lacks negative charge on the other side (Haffke et al., 2019). There are conserved features in binding to SUCNR1, including the interaction with R281^7.39^ and Y83^2.64^, that stabilize the ligands in the suitable positions in both active and inactive SUCNR1 structures. However, the lack of the bulky chemical scaffold in NF-56-EJ40 on the other end adjacent to TM3 predicts a potential role for TM3 in the transition from inactive state to active state (Fig. S12, S13). Upon agonist binding, R99^3.29^ and H103^3.33^ in TM3 move closer to the negatively charged carboxyl group in the ligand, which makes a turning of TM3 while the conserved N105^3.35^ moves away from the hydrophobic site formed by F72^2.53^, F285^7.43^ and L102^3.32^ (Fig. 5F). This helps the rearrangement of these hydrophobic residues upon agonist binding, triggering a π-π stacking between F72^2.53^ and F285^7.43^ (Fig. 5F). The residue rearrangements are transduced through TM3 and TM7. In this case, the side chains of residue such as N287^7.45^, N291^7.49^, Y295^7.53^, Y207^5.58^ and R120^3.50^ serve to break the hydrophobic core formed by F241^6.44^, F245^6.48^, F294^7.52^, I110^3.40^, L113^3.43^, L233^6.36^, L244^6.47^, forcing the outward movement of TM5 and TM6. The hydrogen bond formed by R120^3.50^ and Y207^5.58^ further stabilize the activated conformation (Fig. 5G; Fig. S14). These structural changes are only observed with SUCNR1 in the active state and not in the antagonist-bound structure.

## Discussion

Research on succinate interaction with its receptor SUCNR1 has been hampered in the absence of a high-resolution structure of the agonist-bound receptor. Previous studies used the purinergic receptor P2Y1 for computational modeling based on sequence homology between the two receptors (Li et al., 2023; Trauelsen et al., 2017; Zhang et al., 2015). This approach has inherent limitations because P2Y1 does not bind succinate, and SUCNR1 does not bind any nucleotide. Moreover, P2Y1 is primarily coupled to the Gq class of G proteins whereas SUCNR1 couples primarily to the Gi proteins. The present study addresses these limitations by solving the cryo-EM structures of the succinate-SUCNR1-Gi protein complex and the epoxysuccinate-SUCNR1-Gi protein complex. The two agonist-bound structures represent SUCNR1 in the active state, shedding light on the structural features for G protein activation.

Our structural analysis of the SUCNR1-Gi complex has identified a transmembrane binding pocket that is large relative to the size of succinate (m.w. 118.09). The binding pocket is surrounded by TM1, TM2, TM3 and TM7, and capped by the β-hairpin of ECL2. Since succinate is a dicaboxylate that carries negative charge on both carboxyl groups, positively charged amino acid sidechains are primary suspects for direct interaction with these carboxyl groups (Haffke et al., 2019; Trauelsen et al., 2017). R281^7.39^ is the only positively charged amino acid on one side of the binding pocket with properly placed -NH for interaction with one of the carboxyl groups in succinate. In our structural model, R281^7.39^ forms a salt bridge with this carboxyl group that also interacts with the hydroxyl group of Y30^1.39^ in TM1 through a hydrogen bond. The agonist is further stabilized through an interaction of this carboxyl group with Y83^2.64^, forming another hydrogen bond. R99^3.29^ and H103^3.33^, that were proposed to interact with this carboxyl group in other studies (Geubelle et al., 2017; He et al., 2004), are not found in our model. A water molecule is found in our structural model adjacent to Y83^2.64^. The binding pocket is hydrophilic, but cryo-EM is limited in detecting the presence of water molecules compared to crystallization (Haffke et al., 2019).

There were also differences in the interaction between sidechains of amino acids in the binding pocket and the other carboxyl group of succinate. One of the homology models predicted that R252^6.55^ is next to succinate and interact with its carboxyl group through hydrogen bonding (Geubelle et al., 2017; He et al., 2004). Another paper published in the same year proposed that R255^6.58^, just one turn above R252^6.55^ in TM6, interacts directly with the carboxyl group (Trauelsen et al., 2017). In our molecular model, both R252^6.55^ and R255^6.58^ are too far away from the carboxyl group to allow for the formation of hydrogen bond. In place of arginine in the transmembrane domains, F175^ECL2^ forms a hydrogen bond with the hydroxyl group through the nitrogen in its backbone. F175^ECL2^ is a part of the β-hairpin in ECL2 that folds downward to occlude the binding pocket in the apo state, although a part of the β-hairpin was not visible in the published structure (Haffke et al., 2019). β-hairpins are also observed in P2Y1 and P2Y12 and serve to occlude these receptors (Fig. S15) (Li et al., 2023; Zhang et al., 2015). In our cryo-EM model of active state SUCNR1, the β-hairpin structure is intact and clearly visible. In addition to F175^ECL2^, D174^ECL2^ in the β-hairpin plays a role in stabilizing succinate binding through a salt bridge formed with R281^7.39^. These interactions were not observed in previous computational models and the crystal structure of an antagonist-bound SUCNR1 (Haffke et al., 2019). Our analysis of the cryo-EM structure indicates that the ECL2 β-hairpin interaction with succinate acts as a lid that secures the agonist in the binding pocket. Site-directed mutagenesis of F175^ECL2^ markedly reduced succinate-induced activation of SUCNR1, as evidenced by results from functional assays.

Epoxysuccinate is a cyclic analogue and non-metabolite derivative of succinate (Geubelle et al., 2017). The introduction of the oxygen restricts the rotation of backbone carbon atoms, making this agonist in either *cis* or *trans* conformation. Ligand-receptor-Gi complex was formed only with *cis*-epoxysuccinate, consistent with the finding that succinate is in *cis* conformation when bound to SUCNR1. Within the binding pocket, *cis*-epoxysuccinate interacts through its carboxyl groups with a number of amino acid sidechains that also participate in the binding of succinate, including R281^7.39^ (salt bridge) and Y83^2.64^ (hydrogen bond). Different from succinate, epoxysuccinate binding does not involve Y30^1.39^ and F175^ECL2^; instead, R99^3.29^ forms a salt bridge with the other carboxyl group. This interaction is highly important for the binding of epoxysuccinate as evidenced by functional assay of the site-directed mutant. Two tyrosine residues, Y248^6.51^ and Y277^7.35^, participate in the interaction with the carboxyl group through a water molecule, further securing epoxysuccinate in the binding pocket. These additional interactions, based on our structural model, explain the 10-fold higher potency of epoxysuccinate over succinate at SUCNR1 (Geubelle et al., 2017).

Previous studies using computational modeling and crystallization of antagonist-bound SUCNR1 did not show the receptor-G protein interface. In our structural model, hydrophobic interactions dominate the interface between Gi alpha subunit and SUCNR1. A hydrophobic network consisting of conserved residues A224^6.27^, L225^6.28^, L233^6.36^, L227^6.30^, P230^6.33^, I123^3.53^, P127^34.50ICL2^ and F128^34.51ICL2^ interacts with the α5 helix of Gαi involving I343^G.H5.15^, I344^G.H5.16^, L348^G.H5.20^ and L353^G.H5.25^. There are small differences in the receptor-Gi interface when the succinate-bound and epoxysuccinate-bound SUCNR1 structures are compared. In the succinate-bound SUCNR1 structure, there is a salt bridge between R217^5.68^ and D341 of Gi and a weak hydrogen bond between P127^ICL^ and N347 (Gi) in the backbone. In comparison, the epoxysuccinate-bound SUCNR1 interacts with Gi alpha exclusively through hydrophobic interactions. The difference may cause variations in signal strength and possibly bias in agonism.

The structure of SUCNR1 in active state is compared with the crystal structure of antagonist (NF-56-EJ40)-bound SUCNR1, and distinct features are identified. The agonist bound structure has features of activated GPCRs, including outward shift of TM5 and TM6, and inward movement of TM7. SUCNR1 contains activation components of GPCRs including the DRY, PIF and NPxxY motifs. However, SUCNR1 does not have W^6.48^ in the CWxP motif, which is a conserved component of activated GPCRs. At this position, F^6.48^ replaces tryptophan with a different activation mechanism. Compared with the bound antagonist, which has a much larger chemical backbone, succinate and epoxysuccinate can induce a shift of TM3 through R99^3.29^ that is adjacent to the agonist carrying negatively charged carboxyl group. N105^3.35^, with its hydrophobic sidechain departs from the hydrophobic core of F72^2.53^, F285^7.43^ and L102^3.32^. The resulting rearrangement of space location of amino acids and intermolecular interaction may lead to π-π stacking and associated structural changes transmitting across the membrane through TM3 and TM7. Polar residues on TM3 and TM7, including N287^7.45^, N291^7.49^, Y295^7.53^ and R120^3.50^, turn around to the hydrophobic interface formed by TM5 and TM6. Meanwhile, the polar residues on TM3 and TM7 turn to face the hydrophobic residues on TM5 and TM6, propelling the outward shift of TM5 and TM6. In addition, hydrogen bonding between R120^3.50^ and Y207^5.58^ inside the cell further stabilizes the active conformation of SUCNR1.

The cryo-EM structures of the SUCNR1-Gi complex bound to succinate and epoxysuccinate provide direct evidence for the requirement of the agonists in *cis* conformation and with properly spaced backbone. The negatively charged carboxyl groups interact with the positively charged binding pocket surrounded by amino acid sidechains of TM1, TM2, TM3 and TM7. Of interest, ECL2 also plays an important role in the binding of succinate through F175^ECL2^ that forms a hydrogen bond with one of the carboxyl groups and serves as a lid to cap the binding pocket. D174^ECL2^ interaction with R281^7.39^, which is highly important for ligand binding as evidenced by functional assays of the alanine-substituted mutant, further enhances the interaction between ECL2 and succinate. ECL2 does not contribute to the binding of *cis*- epoxysuccinate, which interacts with the sidechains of Y248^6.51^ and Y277^7.35^ to achieve stable binding. Succinate is a small molecule about the average size of an amino acid, and the transmembrane binding pocket of SUCNR1 is relatively large. This raises the possibility that SUCNR1 may have other ligands. Moreover, the transmembrane binding pocket may be further explored for the identification of antagonists and biased agonists. For instance, our work indicates that some of the amino acids that interact with succinate and epoxysuccinate also interact with NF- 56-EJ40, an antagonist of SUCNR1, providing clues for structural requirement of agonism at SUCNR1. It is hopeful that novel agonists, antagonists and biased agonists may be developed using the structural model of SUCNR1 in active state.

## Author Contributions

A.L. conceived, initiated, and designed the whole project. A.L., Y. L. and W. Z. performed the experiments, analysis the results and prepare the figures. A.L. and R.D.Y. supervised the research and wrote the manuscript with input from all authors.

## Data Availability

The atomic coordinates for the succinate-SUCNR1-Gi complex and the epoxysuccinate-SUCNR1-Gi complex have been deposited in the Protein Data Bank with accession codes 8WOG and 8WP1, respectively. The corresponding EM maps have been deposited in the Electron Microscopy Data Bank with accession codes EMD-37686 and EMD-37707, respectively. All data needed to evaluate the conclusions in the paper are present in the main text or the supplementary materials.

