## Supplemental Materials for "Structural insights into agonist binding and activation of succinate receptor 1"

Supplementary Figs. S1 to S15
Supplementary Table S1

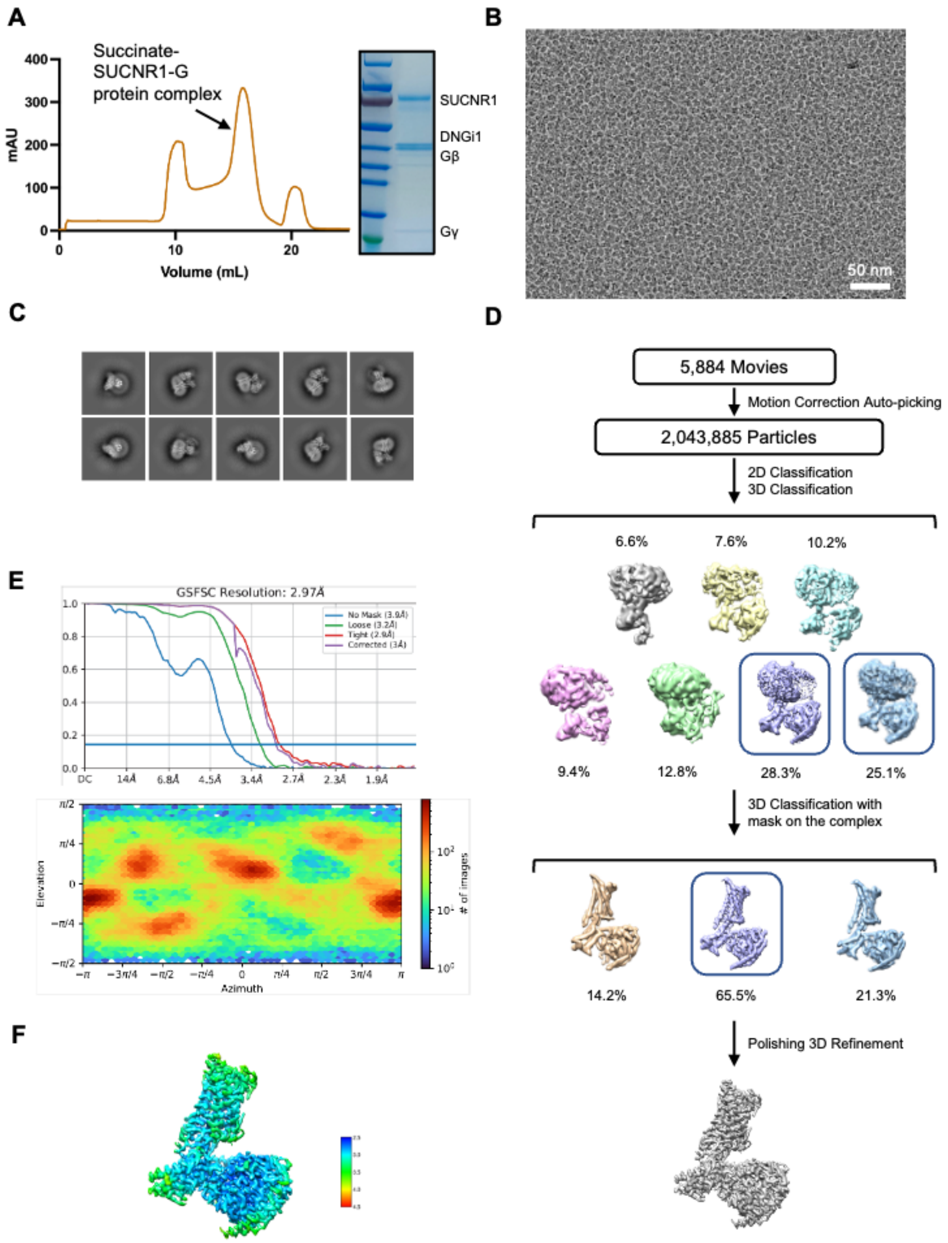


**Supplementary Fig. S1. Purification and cryo-EM data processing of the succiante-bound SUCNR1-Gi complex.**

A. Size-exclusion chromatography profile and SDS-PAGE analysis of the SUCNR1-Gi complex bound with succinate.

B. Representative micrograph after motion correction and dose weighting.

C. 2D class averages of the SUCR1-Gi complex bound with succinate.

D. Workflow of cryo-EM data processing using cryoSPARC.

E. Gold standard Fourier shell correlation (FSC) curve indicates overall nominal resolution at 2.97 Å.

F. Local resolution map.

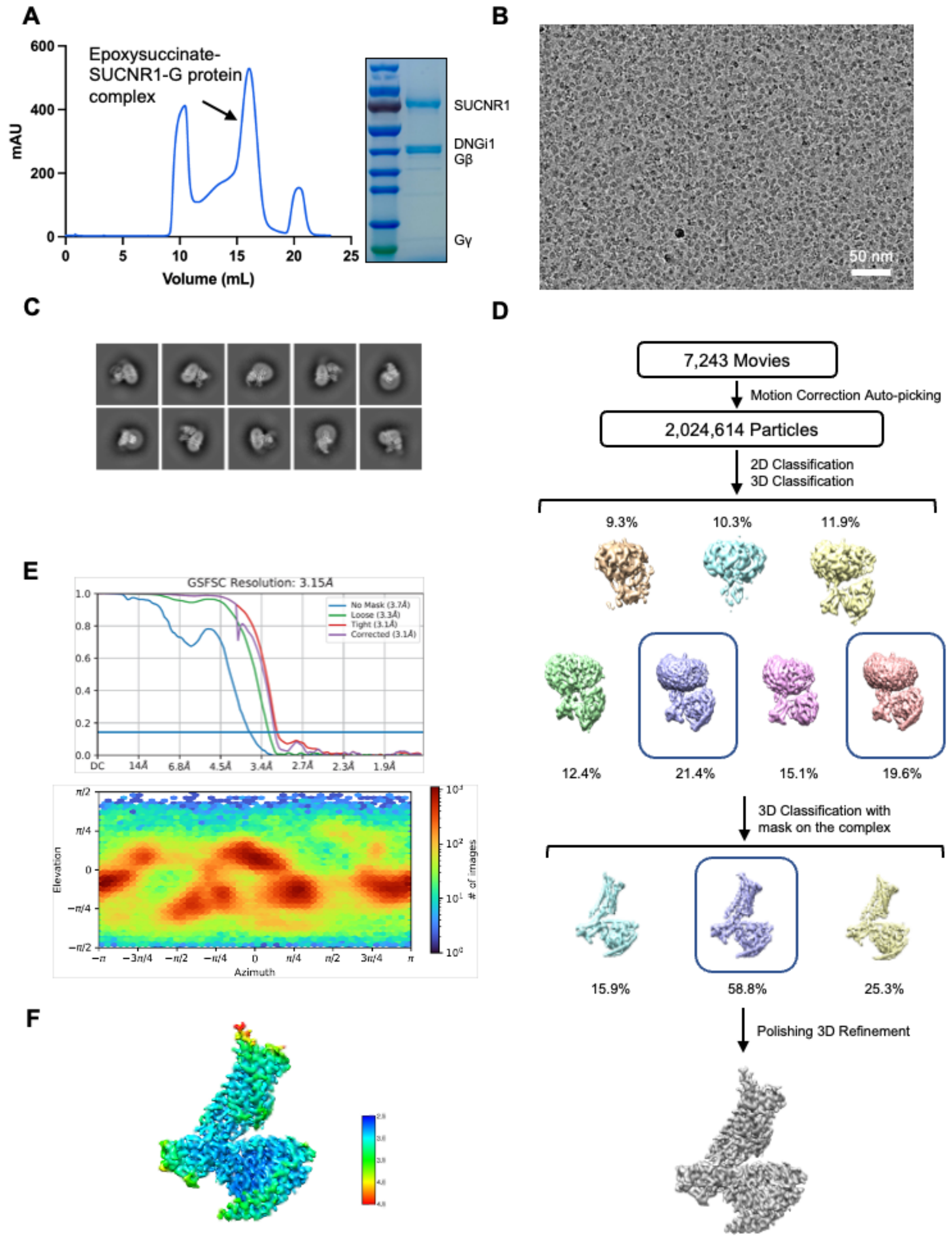


**Supplementary Fig. S2. Purification and cryo-EM data processing of the cis-epoxysucciante-bound SUCNR1-Gi complex.**

A. Size-exclusion chromatography profile and SDS-PAGE analysis of the SUCNR1-Gi complex bound with epoxysuccinate.

B. Representative micrograph after motion correction and dose weighting.

C. 2D class averages of the SUCR1-Gi complex bound with epoxysuccinate.

D. Workflow of cryo-EM data processing using cryoSPARC.

E. Gold standard Fourier shell correlation (FSC) curve indicates overall nominal resolution at 3.15 Å.

F. Local resolution map.

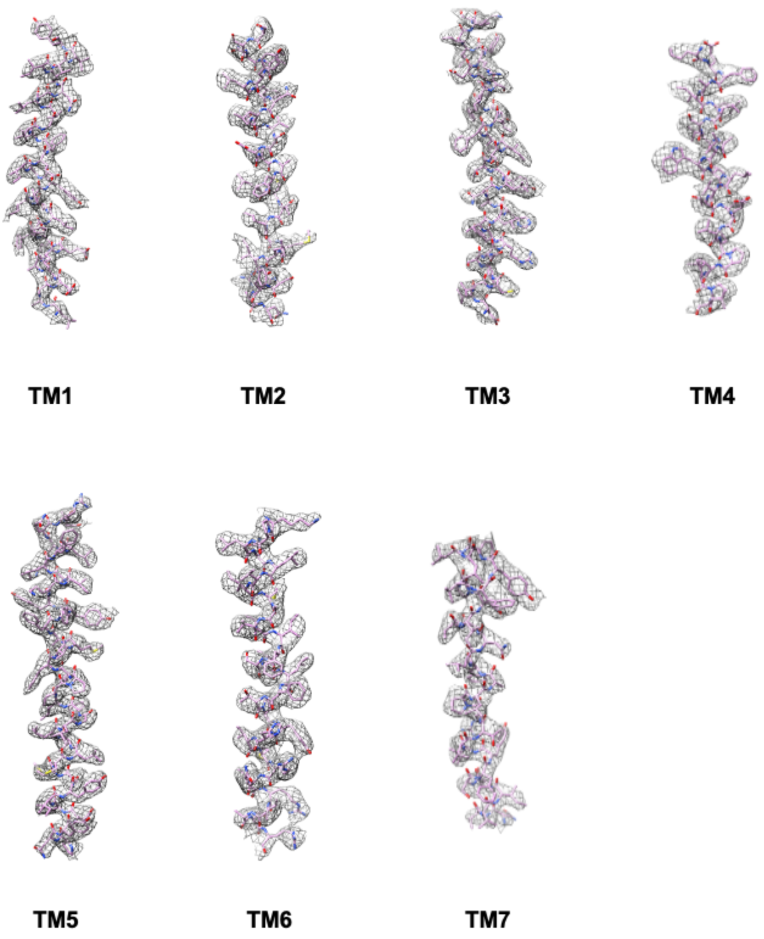


**Supplementary Fig. S3. Representative density maps and models for TM1-7 of succinate-SUCNR1-Gi complex.**

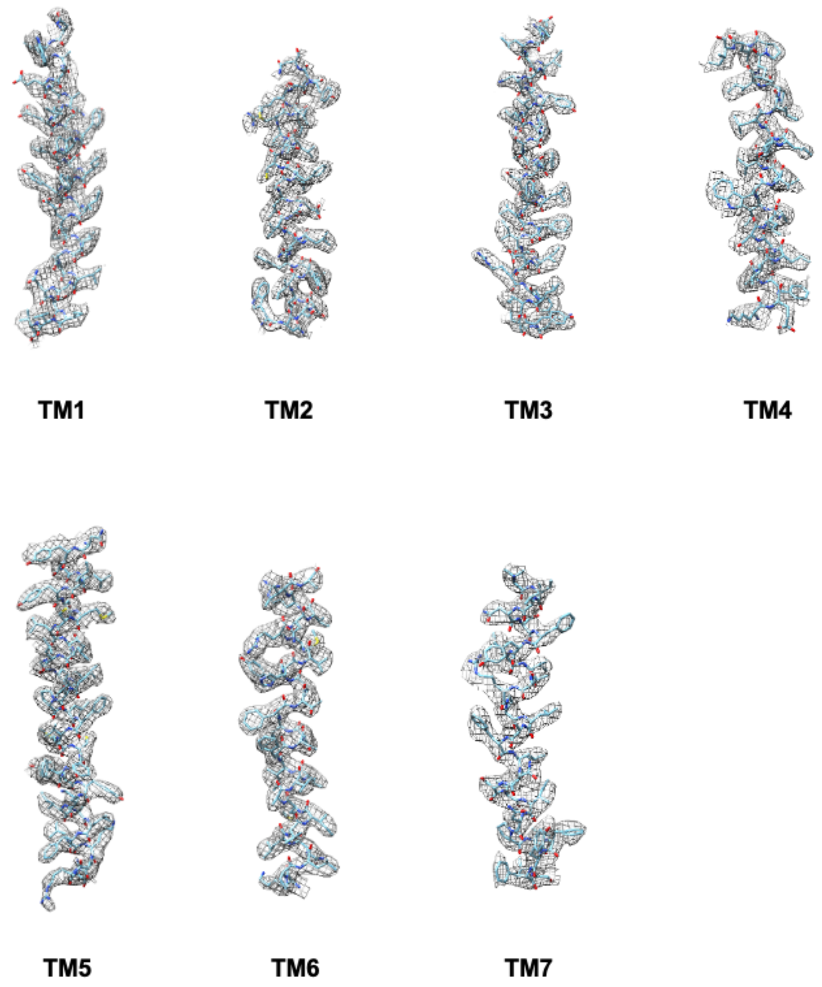


**Supplementary Fig. S4. Representative density maps and models for TM1-7 of epoxysuccinate-SUCNR1-Gi complex.**

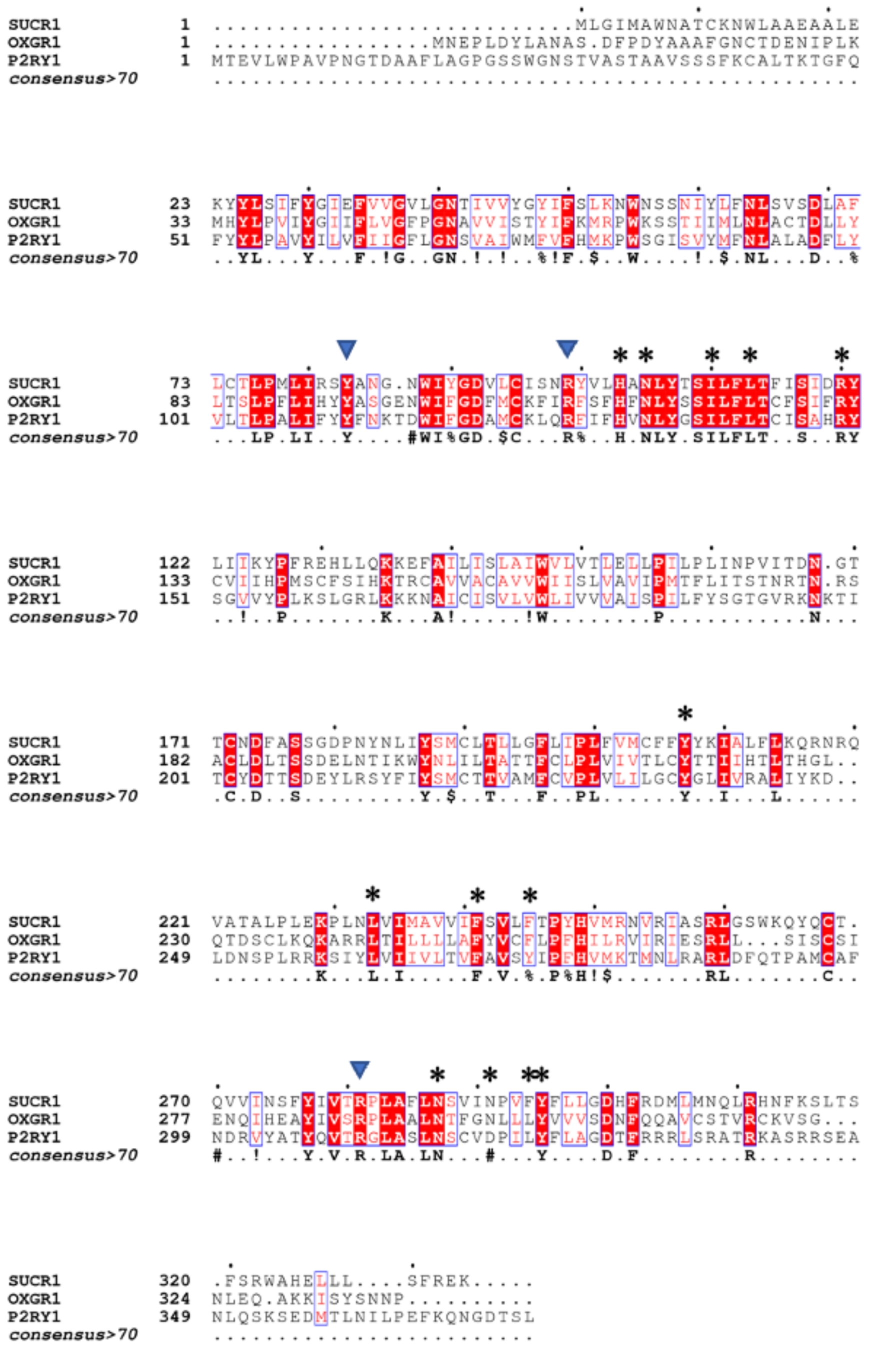

**Supplementary Fig. S5 Sequence alignment of the SUCR1, OXGR1, and P2RY1**. The residues involved in ligand direct binding are marked by blue arrowheads. The asterisk indicates the conserved residue acted in conformational change of transmembrane helix.

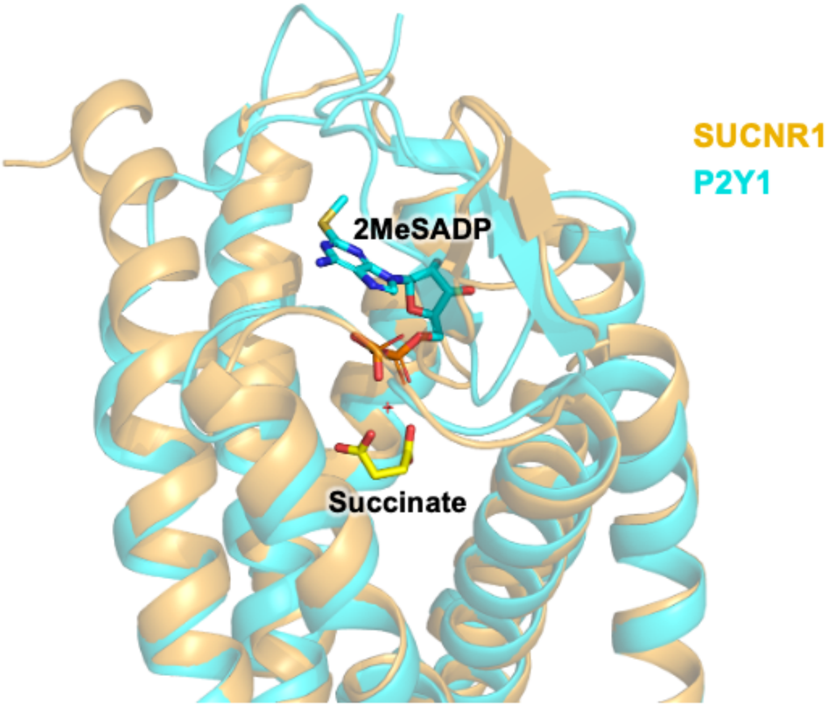


**Supplementary Fig. S6. Structural alignment of the SUCNR1 and the P2RY1 show differences of ligand binding positions.** SUCNR1 structure is shown in orange and succinate is shown in yellow sticks. P2Y1 structure is shown in cyan with the ligand 2MeSADP in cyan sticks.

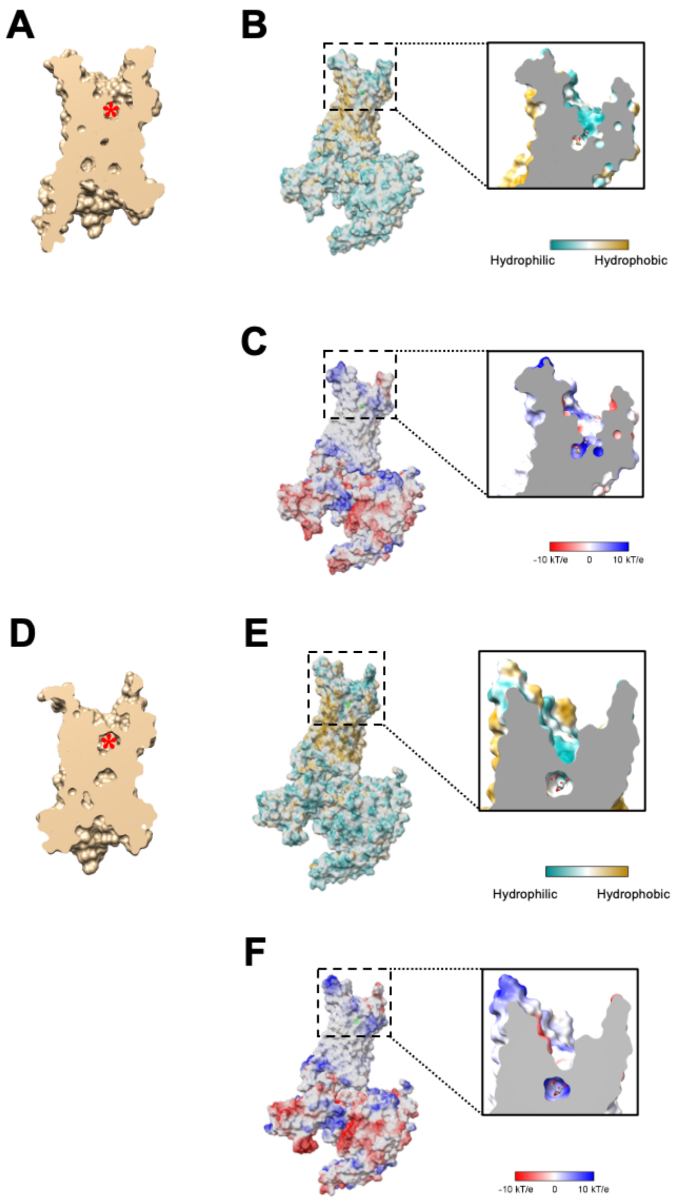


**Supplementary Fig. S7. Slice view of the binding pockets of succinate-bound and *cis*-epoxysuccinate-bound SUCNR1 structure.**
(A) Slice view of the succinate-bound SUCNR1 structure. Ligand position is marked with an asterisk.
(B) Hydrophobicity plot of the succinate binding pocket.
(C) Electrostatic potential plot of the succinate binding pocket.
(D) Slice view of the *cis*-epoxysuccinate-bound SUCNR1 structure. Ligand position is marked with an asterisk.
(E) Hydrophobicity plot of the *cis*-epoxysuccinate binding pocket.
(F) Electrostatic potential plot of the *cis*-epoxysuccinate binding pocket.

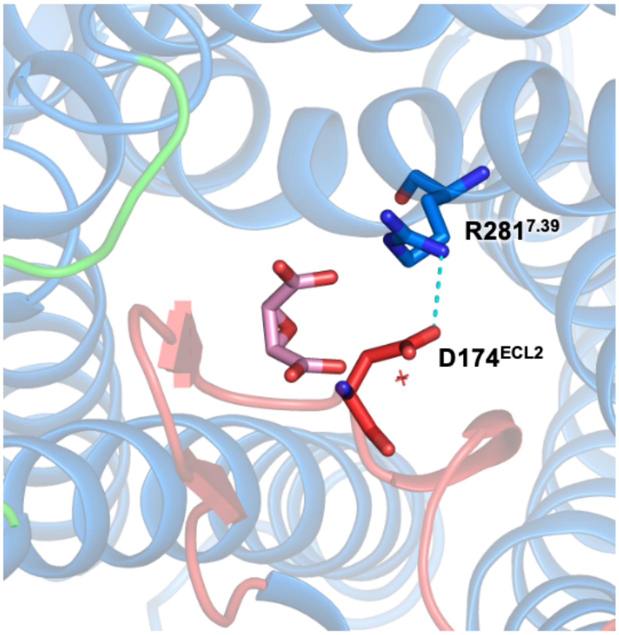


**Supplementary Fig. S8. Top view of the cis-epoxysuccinate binding pocket.** The salt bridge between D174$^{ECL2}$ and R281$^{7.39}$ is shown in cyan dashes.

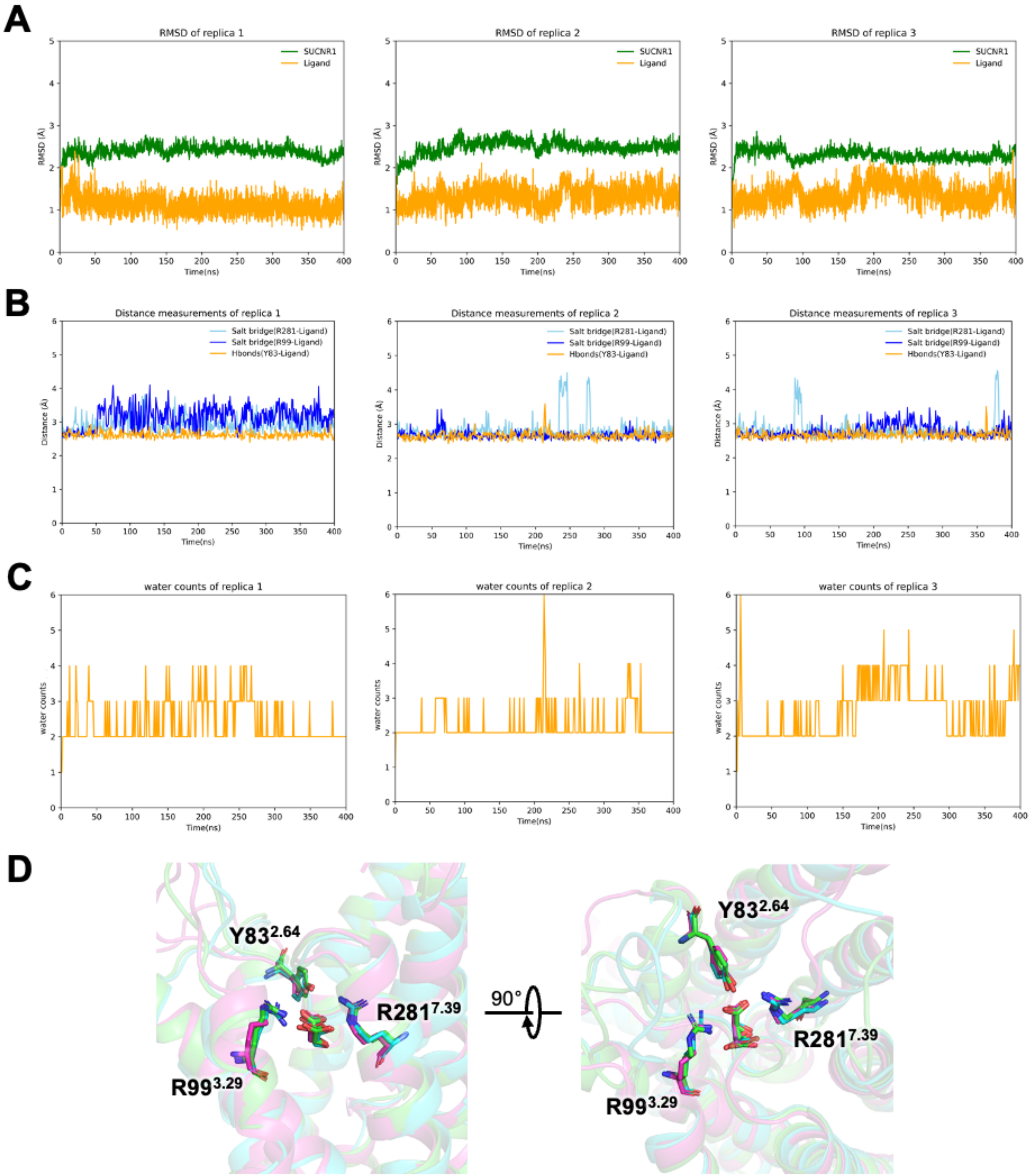


**Supplementary Fig. S9. MD simulation of epoxysuccinate bound SUCNR1.**
(A) RMSD of 3 replicas of epoxysuccinate bound SUCNR1 in 3x400ns MD simulation.
(B) The fluctuations of salt bridge distances and hydrogen bond distances between the ligand and receptor residues R281$^{7.39}$ (salt bridge, cyan), R99$^{3.29}$ (salt bridge, blue) and Y83$^{2.64}$ (hydrogen bond, yellow) in 3x400ns MD simulation.
(C) The variations of water molecule counts near the ligands in a 3.5 Å cutoff in 3x400ns MD simulation.
(D) Representative conformations of the ligand binding pocket in 3x400ns MD simulation.

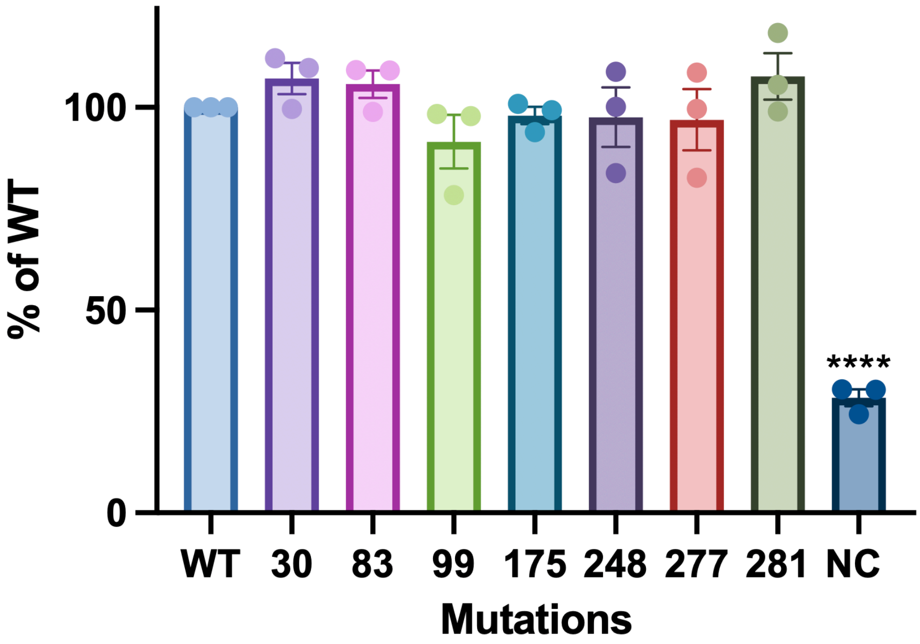


**Supplementary Fig. S10. Surface expression of SUCNR1 mutants.** HEK293T cells were transfected WT or mutant SUCNR1 for 24 h at 37°C. Cells were then incubated with an FITC-conjugated Anti-FLAG antibody for 1 hr on ice. The fluorescence signals on the cell surface were quantified by flow cytometry. WT was set to 100% and NC represents negative control cells transfected with empty plasmids. Data shown are means ± SEM from 3 independent experiments.

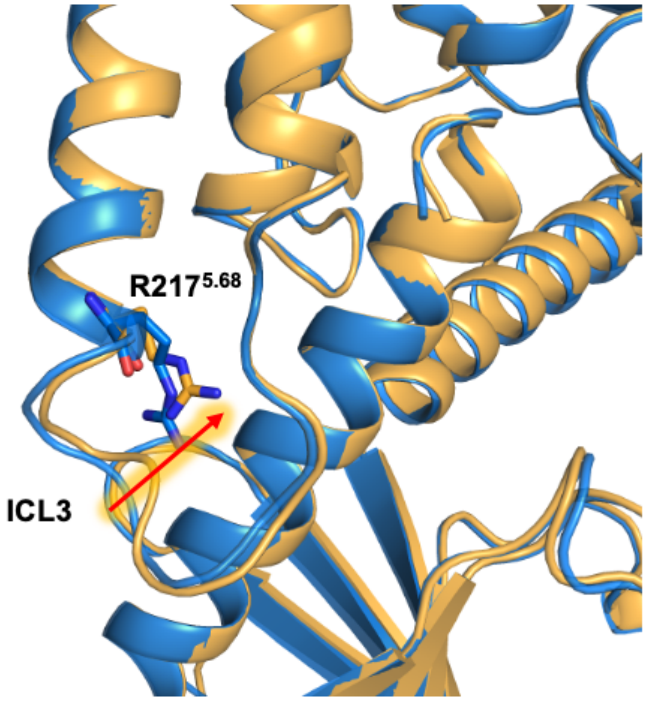


**Supplementary Fig. S11. Subtle differences in ICL3 conformation between succinate-bound and epoxysuccinate-bound SUCNR1.** Inward movements of ICL3 and adjacent R217$^{5.68}$ are highlighted.

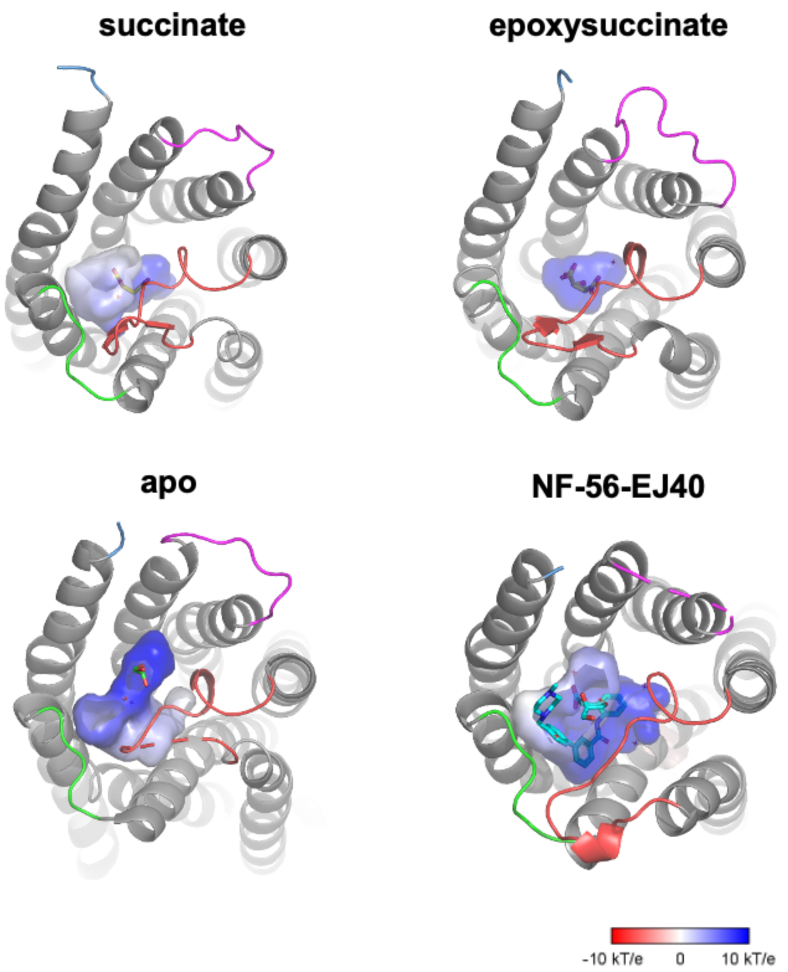


**Supplementary Fig. S12. Ligand binding pockets of succinate-bound, epoxysuccinate-bound, apo and NF-56-EJ40-bound SUCNR1.** Electrostatic potential surfaces for ligand-receptor interactions are shown.

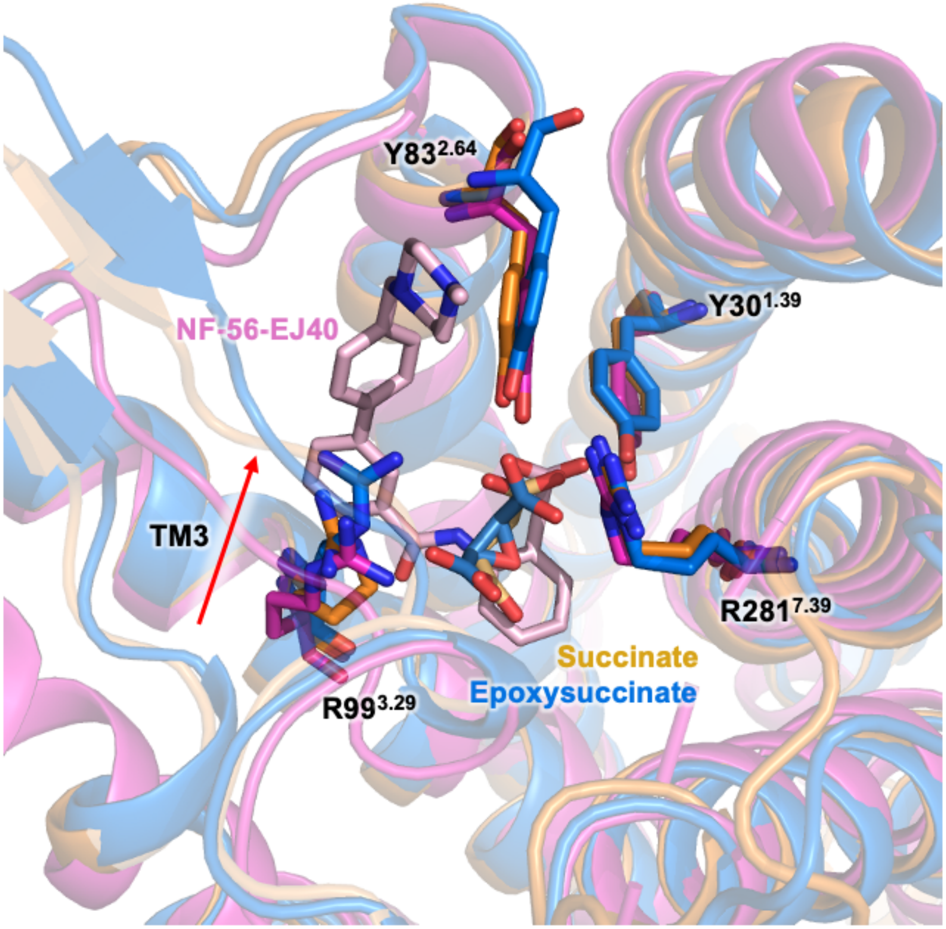


**Supplementary Fig. S13. Comparison of ligand binding pockets between succinate-bound (orange), epoxysuccinate-bound (blue), and NF-56-EJ40-bound (magenta) SUCNR1.** Inward movement of TM3 is highlighted with a red arrow.

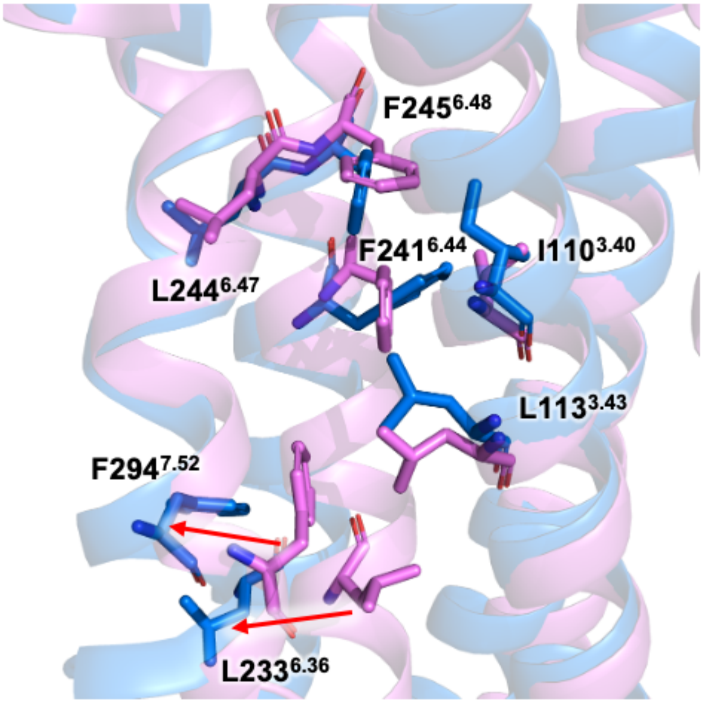


**Supplementary Fig. S14. Hydrophobic network rearrangement at TM3 and TM7.** Movements of TM helices are highlighted in red arrows.

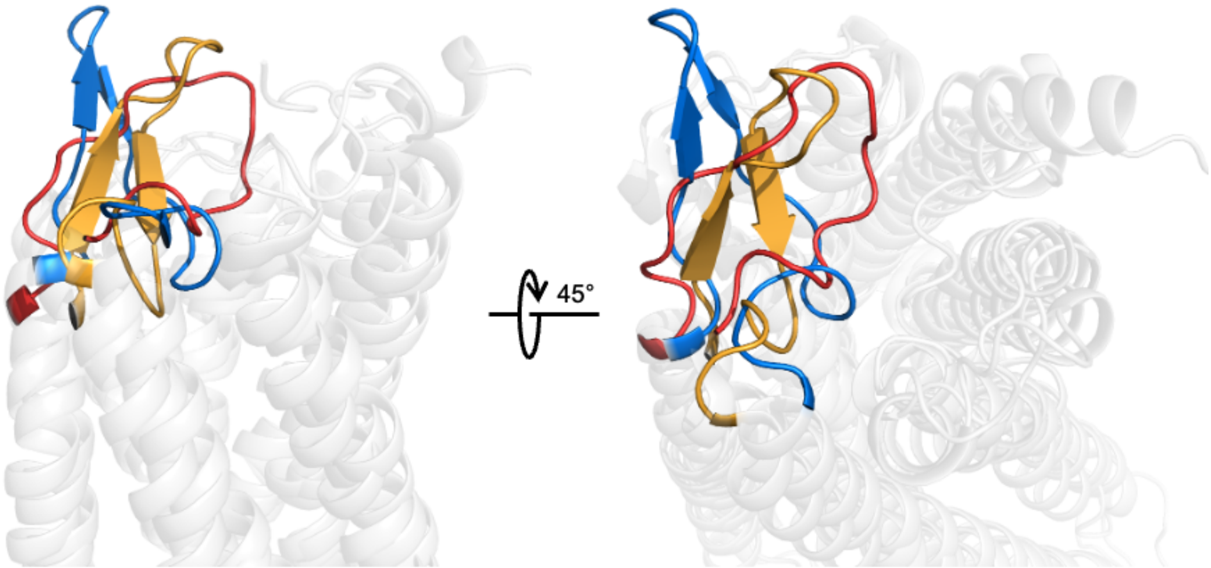


**Supplementary Fig. S15. ECL2 of SUCNR1, P2Y1 and P2Y12 receptors.** ECL2 region of SUCNR1 is shown in blue, ECL2 region of P2Y1 receptor is shown in yellow and ECL2 region of P2Y12 receptor is shown in red. For SUCNR1 and P2Y1 receptor, β-hairpin is observed at ECL2. For all these three receptors, ECL2 occludes the space at the top of the transmembrane binding pocket.

**Supplementary Table S1. Cryo-EM data collection, model refinement and validation statistics.**

| **Data collection and processing** | **Succinate-SUCR1-$G_i$** | ***cis*-epoxysuccinate-SUCR1-$G_i$** |
|---|---|---|
| Magnification | 105000 | 105000 |
| Voltage (kV) | 300 | 300 |
| Electron exposure (e–/Å2) | 52.0 | 52.0 |
| Defocus range (μm) | -1.2~-2.5 | -1.2~-2.5 |
| Pixel size (Å) | 0.85 | 0.85 |
| Symmetry imposed | C1 | C1 |
| Initial particle projections (no.) | 2043885 | 2024614 |
| Final particle projections (no.) | 147371 | 271253 |
| **Refinement** | | |
| Initial model used (PDB code) | 6RNK | 6RNK |
| Map resolution (Å) | 2.97 | 3.15 |
| FSC threshold | 0.143 | 0.143 |
| **Model composition** | | |
| Non-hydrogen atoms | 7382 | 7326 |
| Protein residues | 929 | 922 |
| **R.m.s. deviations** | | |
| Bond lengths (Å) | 0.002 | 0.005 |
| Bond angles (°) | 0.446 | 0.578 |
| **Validation** | | |
| MolProbity score | 2.19 | 2.02 |
| Clashs core | 14.12 | 13.55 |
| **Ramachandran plot** | | |
| Favored (%) | 96.07 | 94.39 |
| Allowed (%) | 3.93 | 5.61 |
| Outliers(%) | 0.00 | 0.00 |
